# Integrating evolutionary and protein language model representations for thermodynamically consistent stability prediction

**DOI:** 10.64898/2026.11.09.765390

**Authors:** Guido Barducci, Tiziana Sanavia, Piero Fariselli

## Abstract

At fixed thermodynamic conditions, protein stability changes obey reversal antisymmetry and path independence, but learned predictors need not satisfy these properties. Explicit alignment-derived evolutionary information may also complement single-sequence protein language model representations, motivating their integration under clearly defined physical constraints.

Here, we develop a training-free strategy that combines thermodynamic constraints with complementary sequence representations. We first introduce EvoPhi-DDG, a potential-based adaptation of the alignment-conditioned evolutionary predictor MSAesm_ddG. By antisymmetrizing its predictions and setting the hidden biases to zero, we express stability changes as differences of a learned, evolution-informed effective potential conditioned on the mutation site and alignment context. This guarantees exact reversal consistency and exact same-site transitivity under fixed alignment contexts while preserving predictive accuracy. We further extend the adapted predictor to multiple substitutions through a background-aware construction that preserves whole-variant antisymmetry.

We then combine EvoPhi-DDG with the previously published JanusDDG using fixed, equal-weight averaging to test predictive complementarity without additional training. The ensemble preserves exact antisymmetry and reduces multisite pathway-closure RMSE relative to JanusDDG across all evaluated path lengths. Furthermore, the ensemble outperforms both components on S669 and S96, achieving Pearson correlations of 0.59 and 0.55, respectively, compared with 0.55 and 0.52 for JanusDDG and 0.53 and 0.48 for EvoPhi-DDG. On the multiple-substitution benchmark PTmul-NR, the ensemble achieves a Pearson correlation of 0.66, improving on JanusDDG’s 0.61 and matching the EvoPhi-DDG extension at the reported precision.

Together, these findings distinguish two benefits: averaging complementary predictions improves accuracy, whereas the potential-based reformulation strengthens physical consistency. They support the integration of alignment-conditioned and single-sequence representations without additional training, while separating predictive improvements from mathematical guarantees.

## 1 Introduction

Protein thermodynamic stability constrains folding, function, molecular evolution and the engineering of useful biomolecules. Amino-acid substitutions can alter the balance between folded and unfolded states, contributing to disease-associated loss of function or enabling the development of more robust proteins. Predicting these effects remains challenging because the relevant free-energy differences are small, sequence context is extensive, and available measurements vary in their experimental conditions and quality. The difficulty increases for multiple substitutions, whose effects need not equal the sum of single-mutant effects. Careful external benchmarking is therefore essential [1, 2].

Computational approaches range from empirical energy functions, exemplified by FoldX, to statistical and machine-learning predictors [3–5]. Evolutionary information provides a complementary perspective: sequence conservation and patterns of tolerated substitutions reflect constraints accumulated across related proteins. Methods such as INPS and DDGun exploit sequence-derived and evolutionary descriptors for stability prediction [4, 6]. These signals are informative but are not synonymous with folding stability, because natural selection also acts on function, expression and molecular interactions.

Protein language models extend this approach by learning contextual representations from large sequence collections. ESM-2 supplies single-sequence embeddings that have been adopted by predictors such as PROSTATA and DDGemb [2, 7, 8]. MSA Transformer instead explicitly conditions on a family alignment through attention across aligned residues and homologous sequences [9]. MSAesm_ddG adapts this representation to experimental stability changes [10].

Both approaches use protein language model representations; the distinction considered here is between explicit family-alignment conditioning and single-sequence inference.

Predictive accuracy alone, however, does not ensure physical consistency. At fixed thermodynamic conditions, the Gibbs free energy is a state function; a change between two protein variants must reverse sign when the variants are exchanged, and must be independent of the path connecting them. Consequently, a stability-change predictor should satisfy two thermodynamic constraints: antisymmetry,

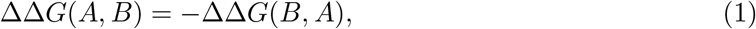

and transitivity,

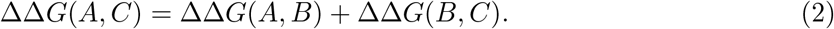

Violations can produce contradictory predictions even when aggregate correlation with experimental labels is high. The importance of reversal consistency has been highlighted in comparative benchmarks [1].

The previously published JanusDDG model combines ESM-2 representations and bidirectional cross-attention with exact antisymmetry at inference and transitivity-oriented post-training [11]. Its sequence-only formulation also supports multiple substitutions. These developments, along-side other thermodynamically regularized sequence models [12], motivate an alternative to training a larger joint model: combine existing predictors with complementary representations while preserving their physical constraints.

We therefore ask whether an existing alignment-conditioned predictor can be reformulated as a site-conditioned potential without retraining, and whether its predictions provide useful information complementary to those of the published JanusDDG model.

Here, we develop EvoPhi-DDG, a thermodynamically constrained, potential-based adaptation of the evolution-informed MSAesm_ddG model that enforces exact antisymmetry, extends prediction to multiple substitutions while preserving this constraint, and guarantees exact same-site transitivity under fixed alignment contexts. We establish the architectural basis of this exact same-site transitivity and use the published JanusDDG model as a fixed ensemble component to investigate predictive complementarity. This combination integrates the contextual single-sequence representations of ESM2 used by the physics-informed JanusDDG model with the explicit evolutionary information provided by family alignments in EvoPhi-DDG. We test accuracy on single and multiple variants, including S669, S96 and PTmul-NR, and evaluate physical consistency separately to distinguish the benefits of prediction averaging from the guarantees introduced by the reformulation.

## 2 Results

### 2.1 EvoPhi-DDG: a potential-based predictor for single and multiple substitutions

To enforce the thermodynamic antisymmetry absent from the original evolution-informed MSAesm_ddG predictor (*M*), we reformulate it as a weight-sharing Siamese model evaluated in both mutation directions:

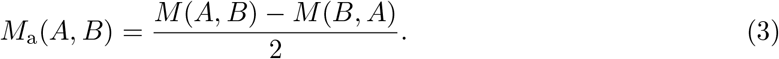

This formulation guarantees exact antisymmetry while retaining the published model parameters and requiring no additional training. Analysis of the resulting head reveals a further opportunity to enforce thermodynamic consistency: setting its hidden biases to zero yields an exact potential-difference formulation, which we term EvoPhi-DDG:

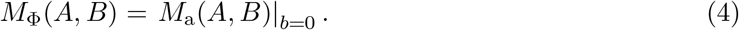

This modification retains the published encoder and weight matrices and guarantees exact same-site transitivity under fixed alignment contexts. Removing the hidden biases produced only minor changes in the predicted stability differences (Supplementary Fig. S1), supporting the adoption of the zero-bias formulation to enforce exact same-site transitivity while closely preserving the original antisymmetric predictions.

We further extend the formulation to multiple substitutions, which are not addressed by the original predictor [10]. We combine single-substitution predictions evaluated in the wild-type and fully mutated sequence backgrounds, followed by antisymmetrization of the complete variant prediction. This background-aware construction preserves exact antisymmetry for multiple substitutions without additional training (see Methods, Section 3.4). Figure 1 illustrates the EvoPhi-DDG formulation and its extension to multiple substitutions while preserving whole-variant antisymmetry.

**Figure 1:**
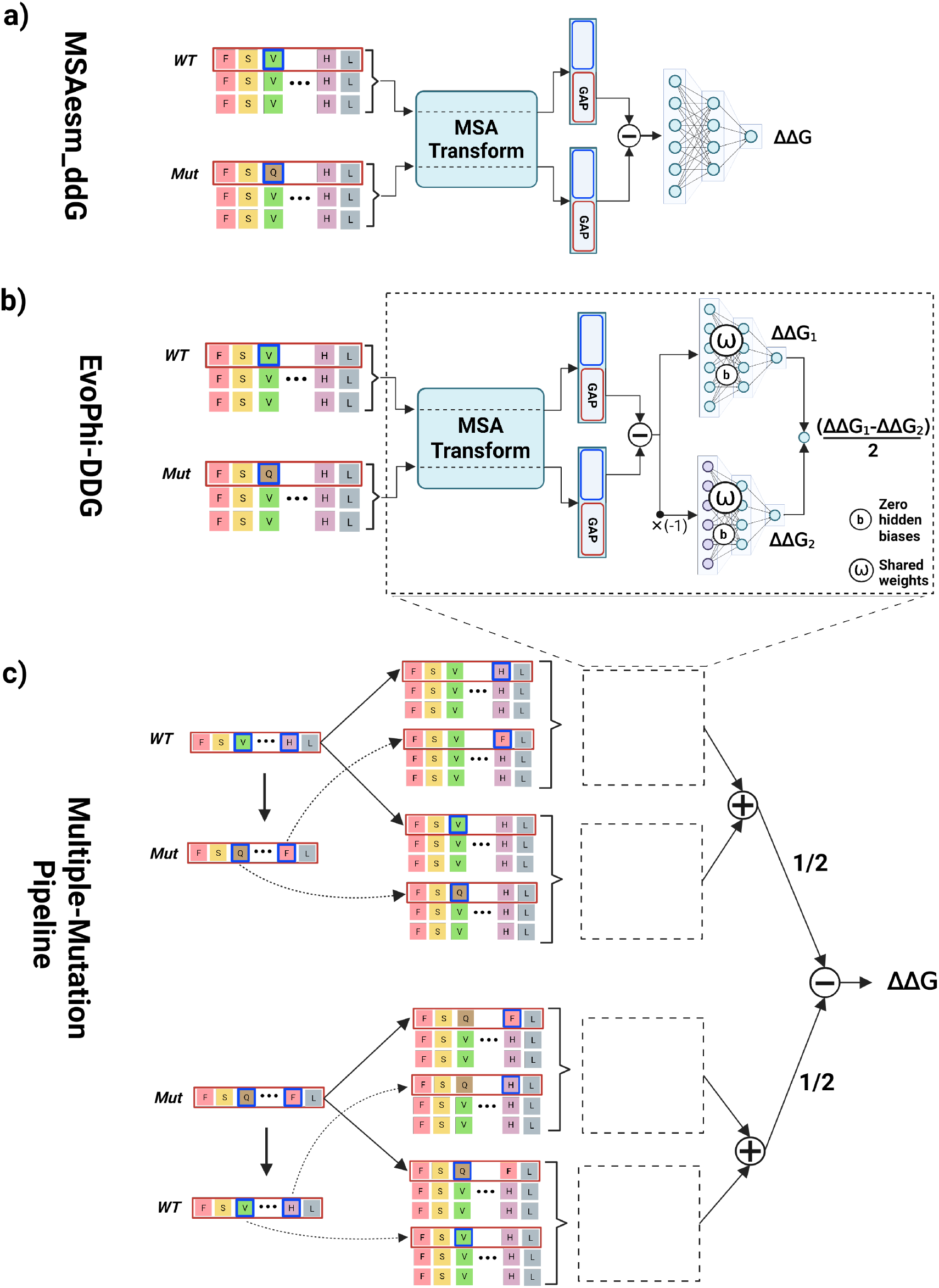
EvoPhi-DDG formulation and extension to multiple substitutions. **a**, The original MSAesm_ddG model produces a directional stability-change prediction from the representations of the WT and mutant sequences. **b**, EvoPhi-DDG evaluates the model with shared parameters and zero hidden biases in both input orientations, then takes half the difference between the directional outputs. This yields a potential-difference formulation that guarantees exact antisymmetry and exact same-site transitivity under fixed alignment contexts, without additional training. **c**, Extension to multiple substitutions. Individual substitutions are scored independently on the WT background, and their reversions are scored independently on the fully mutated background. The difference between these two sums, divided by two, yields a whole-variant prediction that preserves exact antisymmetry.

In this section, we assess the resulting model’s compliance with the thermodynamic constraints of antisymmetry and transitivity, and evaluate its predictive performance relative to the original MSAesm_ddG formulation.

#### 2.1.1 Thermodynamic consistency of EvoPhi-DDG

We first evaluated the two thermodynamic properties, antisymmetry and transitivity, in the original MSAesm_ddG predictor and its antisymmetric Siamese formulation using identical sequence states and alignment contexts. The original model exhibited a mean absolute reversal defect |*M* (*A, B*) + *M* (*B, A*)| of 0.20 kcal/mol, with RMSE 0.28 kcal/mol across S669. The largest defect was 1.19 kcal/mol. Its identity prediction, *M* (*A, A*), was −1.38 × 10^−3^ kcal/mol rather than zero. The antisymmetric Siamese formulation eliminated both defects exactly, as expected from its construction (Methods, Section 3.2.3).

To assess same-site transitivity, we followed the evaluation procedure used for JanusDDG [11], connecting each S669 endpoint pair through 1, 3, 5, 7 or 9 random intermediate residues at the same mutation site, with ten random repetitions for each intermediate count. Alignment rows, windows and the mutation position were held fixed along a path. The original MSAesm_ddG predictor was not transitive: its mean direct-versus-path Pearson correlation decreased from 0.96 with one intermediate to 0.84 with nine, while closure RMSE increased from 0.26 to 0.61 kcal/mol. Antisymmetrization almost eliminated this deterioration: with nine intermediate states, the mean direct-versus-path Pearson correlation was *r* = 1 − 1.38 × 10^−6^, while the mean closure RMSE was 1.50 × 10^−3^ kcal/mol, indicating near-exact but not perfect same-site transitivity (Fig. 2; Supplementary Table S7).

**Figure 2:**
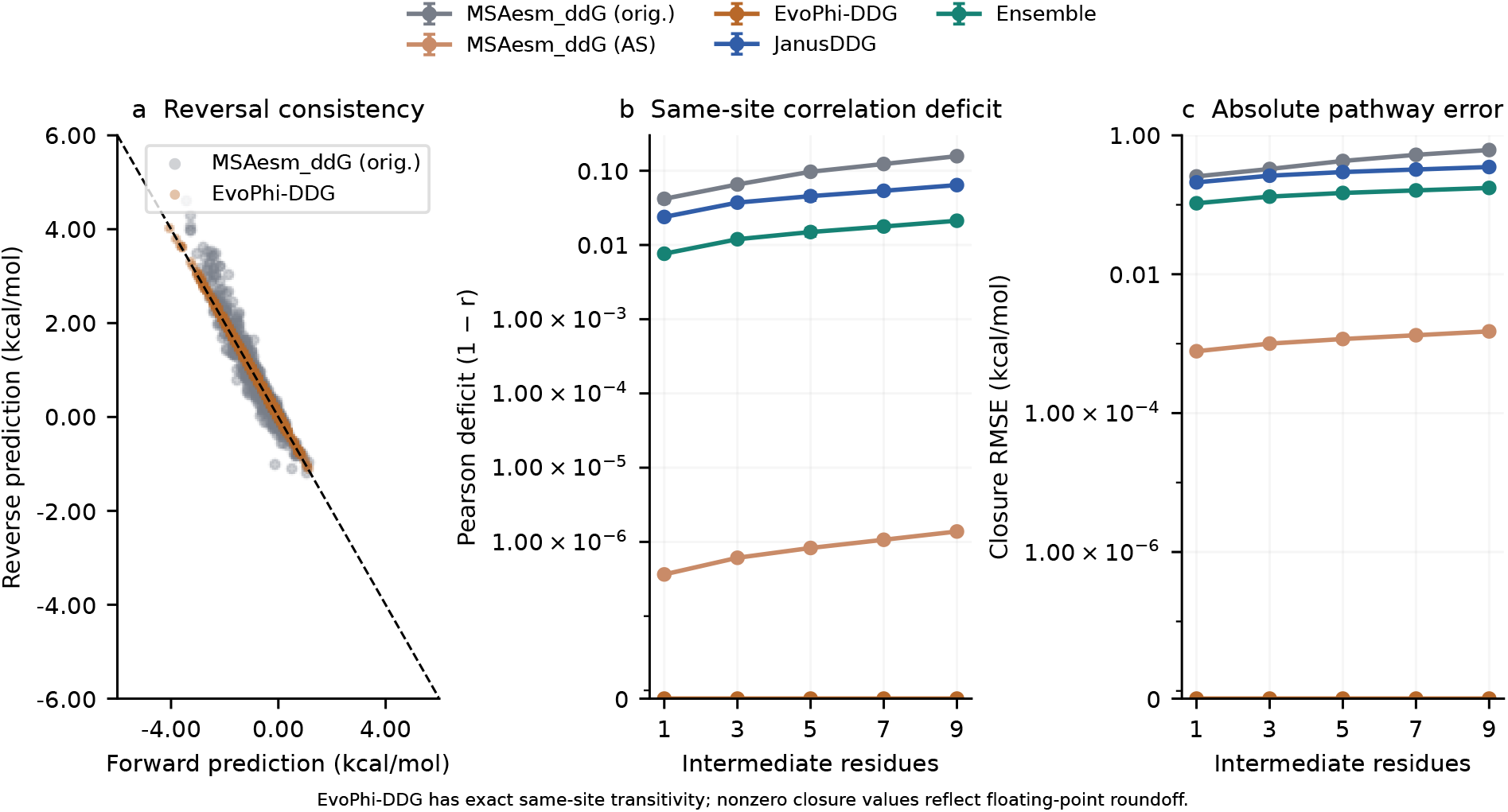
Physical consistency of EvoPhi-DDG and its component controls. **a**, Forward and reverse MSAesm_ddG predictions on S669; the dashed line denotes exact antisymmetry. **b**, Pearson deficit 1 −*r* between direct and pathway-summed predictions on a symmetric logarithmic scale with a linear region around zero. Error bars show the standard deviation over ten random paths per intermediate count. **c**, Closure RMSE on the same type of symmetric logarithmic scale. The linear regions extend to 10^−7^, with a true zero tick; nonzero numerical residuals are retained. Both tests use identical states and paths for all models. These are prediction-versus-prediction consistency measures, not accuracy against experimental labels. The antisymmetrized MSAesm_ddG control retains the published hidden biases; EvoPhi-DDG removes them. Its nonzero numerical closure residuals reflect floating-point precision.

This pronounced improvement has a specific architectural explanation. MSAesm_ddG applies a single-hidden-layer ReLU head to a difference of state representations. If the hidden bias is zero, the identity ReLU(*x*) − ReLU(−*x*) = *x* reduces the antisymmetric head to a linear functional of that difference, yielding an exact potential difference and therefore exact transitivity through cancellation of intermediate-state contributions. For the published nonzero biases, we derive an explicit residual and a uniform bound (Methods, Section 3.2.4).

Together with the observation that setting the hidden biases to zero leaves predictions nearly unchanged (Fig. S1), this analysis motivates the zero-hidden-bias formulation used in EvoPhi-DDG (see Section 3.2.1 for details on the notation):

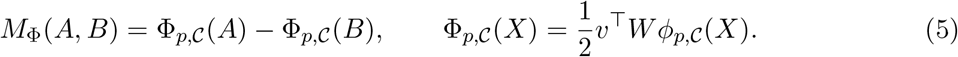

Removing the hidden biases eliminates the nonlinear residual and replaces near-exact same-site transitivity with an exact algebraic guarantee, while preserving predictive performance. The potential is conditioned on the mutation position *p* and alignment context C; this result therefore does not establish general multisite path independence and is not a general consequence of antisymmetry for arbitrary neural architectures.

Given the exact same-site transitivity of EvoPhi-DDG, we next assessed whether this consistency extended to paths involving temporary substitutions at other positions, using the same 669 S669 endpoint pairs. For each WT sequence *A* and its single-substitution mutant *B*, we introduced *m* temporary substitutions at distinct non-target positions, applied the target substitution, and then reverted the temporary substitutions in reverse order:

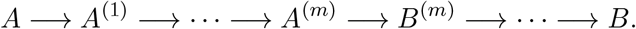

Here, *A*^(*j*)^ and *B*^(*j*)^ carry the first *j* temporary substitutions, without and with the target substitution, respectively. Each edge is a single substitution. The path contains *N* = 2*m* intermediate sequence states and *N* + 1 edges; its endpoints remain the original S669 WT–mutant pair.

We evaluated *N* = 2, 4, 6, 8, 10 intermediates with ten random repetitions per variant and path length, obtaining 33,450 paths across 87 distinct WT proteins. Temporary sites were sampled within each variant’s existing MSA window, and homolog rows, row order, depth and window coordinates were held fixed. The analysis encoded 196,196 context-specific sequence states and scored 229,558 distinct directed single-substitution edges. All predictors used identical paths, with full-length sequences supplied to JanusDDG (Methods, Section 3.5.2).

For original MSAesm_ddG, mean direct-versus-path Pearson correlation decreased from 0.93 with two intermediates to 0.70 with ten, while mean closure RMSE increased from 0.33 to 0.88 kcal/mol. EvoPhi-DDG reduced the corresponding closure RMSEs to 0.04 and 0.11 kcal/mol, with correlations of 1−1.08×10^−3^ and 1−7.21×10^−3^, respectively. At ten intermediates, its mean closure MAE was 0.06 kcal/mol and maximum absolute defect was 1.37 kcal/mol (Supplementary Table S8). The antisymmetric formulation retaining the published hidden biases closely matched EvoPhi-DDG, with the same correlations, mean closure MAEs and RMSEs at the displayed precision.

Thus, eliminating the hidden biases establishes exact same-site transitivity without materially altering multisite consistency. The remaining multisite defects reflect the site-dependent readout, which prevents the fixed-site potential from defining a common potential across different mutation positions.

#### 2.1.2 Predictive performance of EvoPhi-DDG

Predictive accuracy is as essential as thermodynamic consistency for a useful ΔΔ*G* predictor. We therefore assessed whether the improved physical consistency of the potential-based formulation was accompanied by changes in accuracy against experimental measurements.

In terms of predictive performance, antisymmetrization largely preserved the accuracy of MSAesm_ddG. On the primary S669 labels, Pearson correlation changed from 0.54 to 0.53 (Supplementary Table S1), whereas on S96 Pearson correlation increased from 0.47 to 0.48 (Supplementary Table S3). These modest changes indicate no consistent gain or loss in predictive accuracy.

Setting the hidden biases to zero preserved these predictive results. Predictive performance was also preserved for multiple substitutions: on the 82 PTmul-NR variants, EvoPhi-DDG retained a Pearson correlation of 0.66, MAE of 1.45 kcal/mol and RMSE of 1.95 kcal/mol, matching the antisymmetric formulation with the published hidden biases at two-decimal precision (Supplementary Table S4). EvoPhi-DDG therefore retains the predictive performance of the antisymmetric formulation while guaranteeing exact reversal consistency and exact same-site transitivity under fixed alignment contexts.

### 2.2 Original JanusDDG as a fixed reference and ensemble component

We use the original JanusDDG checkpoint without modifying its architecture or weights [11]; the following evaluations provide a common reference for assessing complementarity and physical consistency.

The physics-informed, ESM2-based predictor JanusDDG exhibited exact antisymmetry and strong same-site transitivity, with direct-versus-path correlations ranging from 0.98 to 0.94 as the number of intermediates increased (Fig. 2; Supplementary Table S7). For S669 multisite pathways with ten intermediates, JanusDDG achieved a mean direct-versus-path Pearson correlation of 0.98 and mean closure RMSE of 0.16 kcal/mol (Supplementary Table S8).

Regarding predictive performance, JanusDDG achieved Pearson correlations of 0.55 on S669, 0.69 on its curated subset S461, and 0.52 on S96 for single substitutions. For multiple substitutions, it achieved a Pearson correlation of 0.61 on PTmul-NR (Supplementary Table S9).

### 2.3 Integrating alignment-conditioned and single-sequence predictions

We combined EvoPhi-DDG with the previously published JanusDDG predictor to test whether alignment-conditioned and single-sequence predictions provide complementary information while retaining exact reversal antisymmetry. The ensemble averages the outputs of the two predictors with equal weights, without additional training or calibration (Methods, Section 3.3). A schematic representation of the ensemble model is shown in Fig. 3.

**Figure 3:**
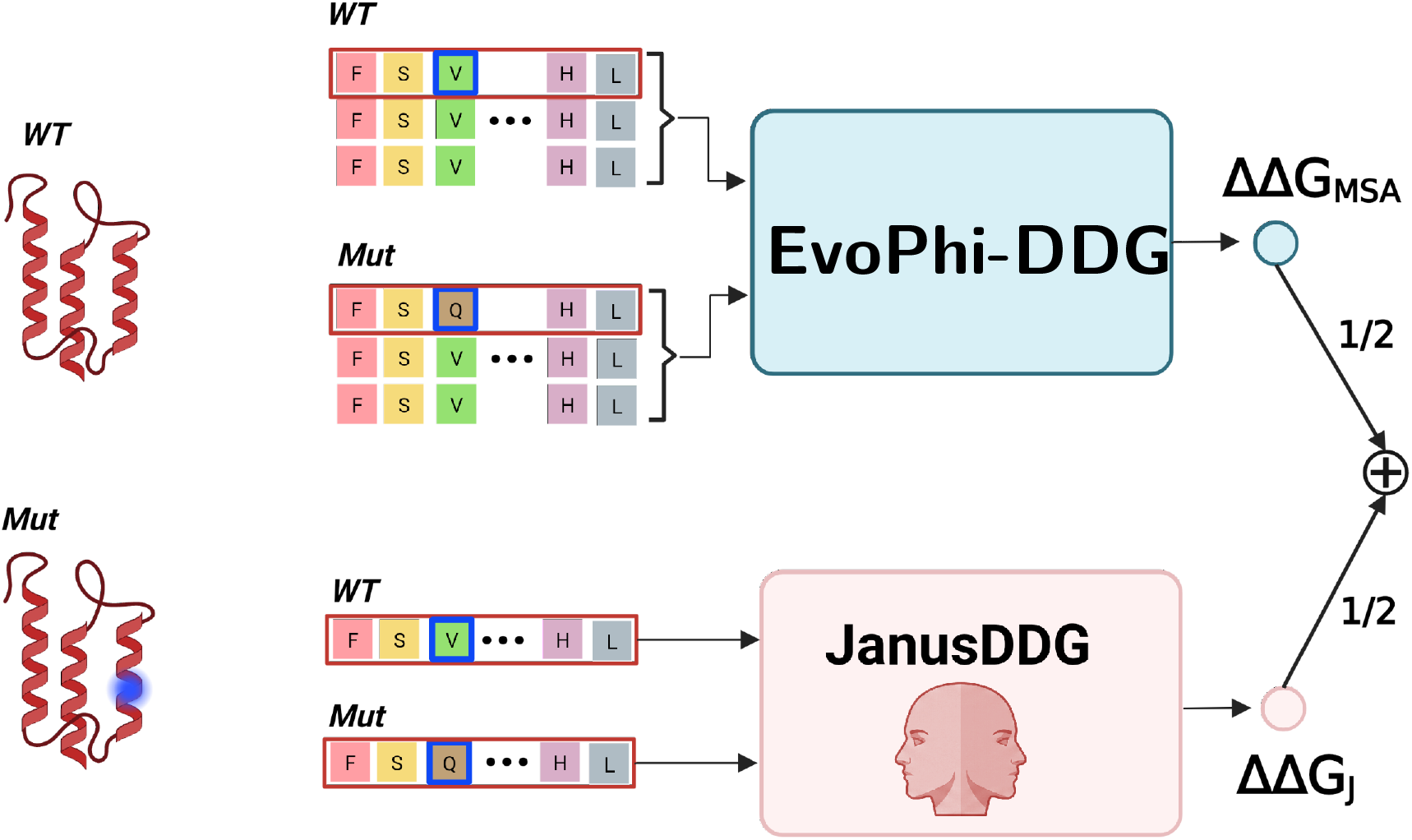
Integration of physics-informed and evolution-informed predictions. The ensemble combines JanusDDG, a physics-informed predictor based on ESM2 sequence representations, with EvoPhi-DDG, a potential-based reformulation of MSAesm_ddG that incorporates evolutionary information from homologous sequence alignments. The two predictions are combined with equal weights, without additional training or calibration. For multiple substitutions, EvoPhi-DDG uses the background-aware extension described in Fig. 1. The resulting ensemble supports single and multiple substitutions while preserving exact antisymmetry.

We evaluate predictive accuracy and physical consistency separately, using the original-head controls in Section 2.5 to distinguish the effects of averaging from those of imposing physical constraints.

#### 2.3.1 Thermodynamic consistency of the ensemble model

The ensemble preserves exact antisymmetry by construction and exhibits strong same-site transitivity. Correlations between direct and pathway-summed predictions were 0.99 and 0.98 for paths with one and nine intermediates, respectively. At nine intermediates, closure RMSE was 0.17 kcal/mol, exactly half that of JanusDDG (0.35 kcal/mol) before rounding (Fig. 2; Supplementary Table S7). This reduction follows from the ensemble’s signed closure error being the mean of the component errors: under fixed mutation-site and alignment contexts, EvoPhi-DDG contributes zero closure error, halving the discrepancy retained by JanusDDG (Methods, Section 3.3).

The improvement extended to multisite pathways on S669. At ten intermediates, the ensemble achieved mean closure RMSE of 0.10 kcal/mol, compared with 0.16 for JanusDDG and 0.11 for EvoPhi-DDG, corresponding to reductions of 37.81% and 10.03%, respectively (Fig. 4; Supplementary Table S8). Its mean direct-versus-path Pearson correlation was 0.99 and mean closure MAE was 0.05 kcal/mol. The ensemble reduced closure RMSE relative to JanusDDG at all five path lengths and relative to EvoPhi-DDG at four of five; with eight intermediates, EvoPhi-DDG had lower closure RMSE (0.09 versus 0.10 kcal/mol). At ten intermediates, within-protein RMSE improved over JanusDDG in 68 of 87 WT groups. The mean RMSE reduction relative to Janus-DDG was 0.06 kcal/mol, with a descriptive 95% paired protein-bootstrap interval of [0.02, 0.10]; the reduction relative to EvoPhi-DDG was 0.01 kcal/mol, with interval [−0.03, 0.04]. Thus, the latter point-estimate advantage was not resolved by the bootstrap interval.

**Figure 4:**
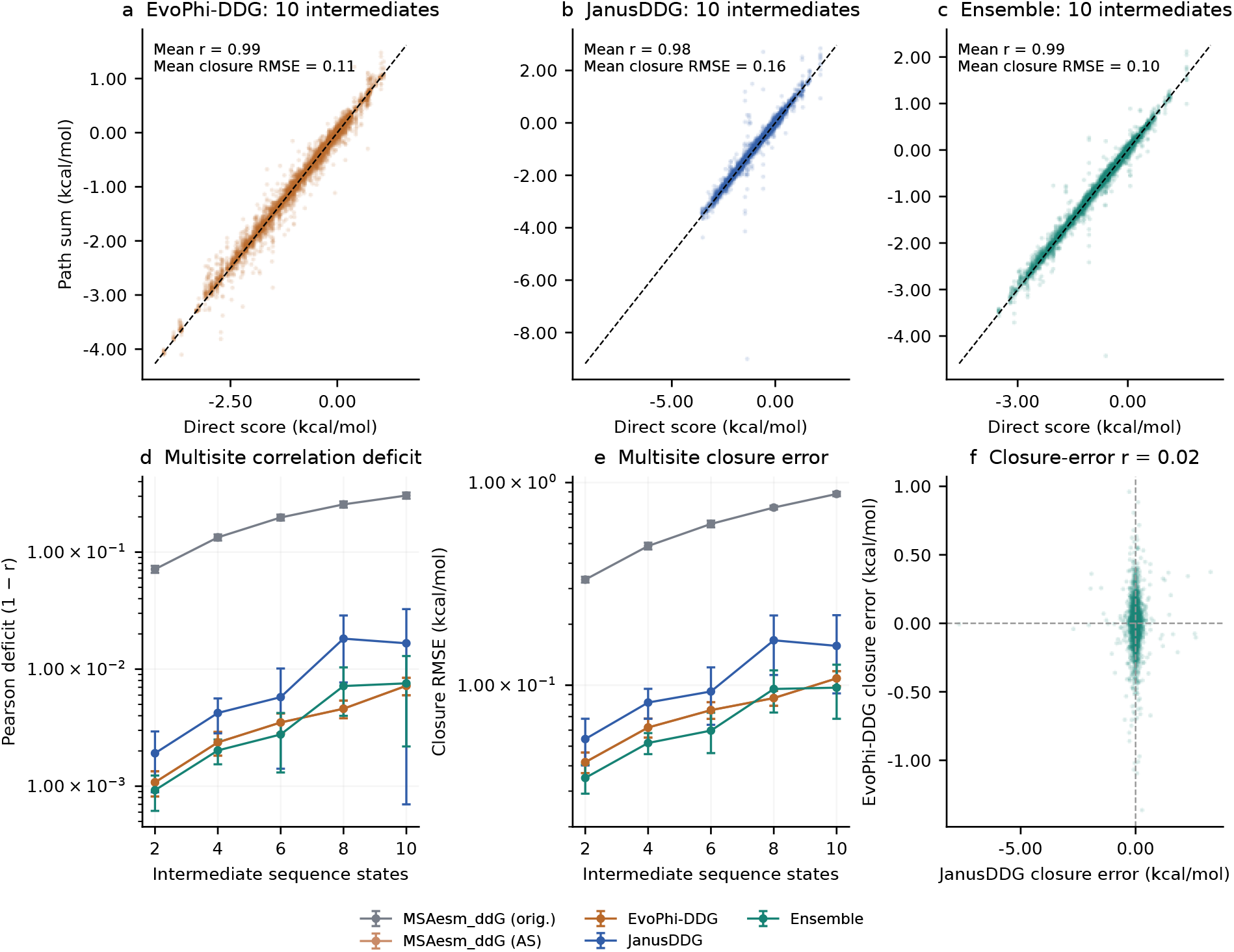
Multisite pathway consistency on S669 without intermediate labels. **a–c**, Direct predictions versus sums along paths with ten intermediate sequence states for EvoPhi-DDG, JanusDDG and their fixed equal-weight ensemble. Each panel shows all 6,690 variant– repetition pairs; annotations report mean Pearson correlation and mean closure RMSE over ten repetitions. Dashed lines indicate identity. **d**, Direct-versus-path Pearson deficit 1 − *r* and **e**, closure RMSE for 2, 4, 6, 8 and 10 intermediates, including the original and antisymmetrized MSAesm_ddG controls. Error bars show standard deviations over ten repetitions; both panels use logarithmic vertical scales. **f**, Signed component closure errors at ten intermediates, across all variant–repetition pairs. Each path temporarily changes non-target sites, applies the target substitution and restores the temporary sites. All comparisons share the same 669 endpoint pairs and paths. The full analysis comprises 33,450 paths on 87 WT-protein groups; paths sharing endpoints or proteins are not independent experimental measurements.

#### 2.3.2 Predictive complementarity on single- and multiple-substitution benchmarks

The equal-weight combination reached Pearson 0.59 and Spearman 0.61 on the 669 direct S669 variants, compared with Pearson 0.55 for JanusDDG and 0.53 for EvoPhi-DDG. MAE decreased to 0.91 kcal/mol and RMSE to 1.33 kcal/mol, improving on both components (Fig. 5a–c; Supplementary Table S9).

**Figure 5:**
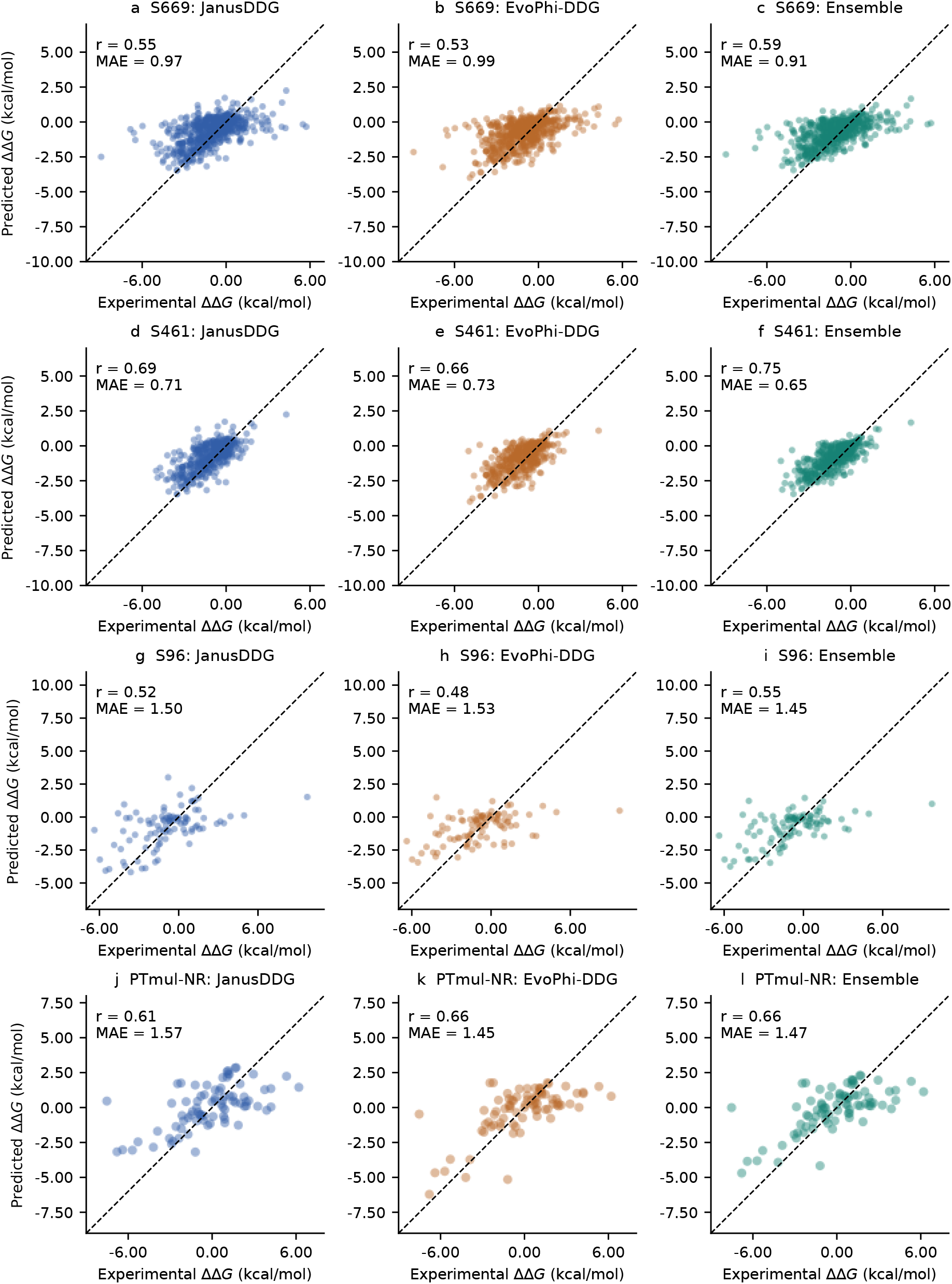
External single- and multiple-mutant prediction. Experimental versus predicted stability changes for S669, its curated S461 subset, S96 and PTmul-NR (top to bottom). Columns show JanusDDG, EvoPhi-DDG and their fixed mean. The EvoPhi-DDG predictions for PTmul-NR use the background-aware, whole-variant-antisymmetric construction. Dashed lines indicate identity. Axes cover all experimental and predicted values; statistics use all variants.

Paired resampling of the 87 distinct full-length WT sequence groups supported a positive improvement on this dataset. The ensemble’s Pearson increase over JanusDDG was 0.05, with a descriptive 95% protein-bootstrap interval of [0.02, 0.08]; the increase over EvoPhi-DDG was 0.06 [0.03, 0.09] (Figure S3, Supplementary Table S6).

Table 1 places the results alongside representative published stability predictors. Importantly, direct-variant scores must not be confused with scores pooled across artificially generated direct and reverse variants. For example, DDGemb reports Pearson 0.53 on direct S669 variants and 0.68 on the pooled set [2]. The ensemble also improved performance on S461 and S96 (Fig. 5, S2, S3).

**Table 1:** Context against published single-mutant benchmarks. Literature values refer to the direct S669 set. Values below the rule are recalculated in this study.

| Method | Source | Pearson | MAE | RMSE |
|---|---|---|---|---|
| DDGemb | [2] | 0.53 | 0.99 | 1.40 |
| MSAesm_ddG, original report | [10] | 0.53 | 0.99 | 1.41 |
| PROSTATA | [2, 8] | 0.49 | 1.00 | 1.45 |
| ACDC-NN | [1, 2] | 0.46 | 1.05 | 1.49 |
| INPS-Seq | [1, 2] | 0.43 | 1.09 | 1.53 |
| ThermoMPNN | [5, 10] | 0.43 | — | 1.52 |
| JanusDDG | This study | 0.55 | 0.97 | 1.39 |
| MSAesm_ddG, current inputs | This study | 0.54 | 0.97 | 1.38 |
| MSAesm_ddG, antisymmetrized | This study | 0.53 | 0.99 | 1.39 |
| EvoPhi-DDG | This study | 0.53 | 0.99 | 1.39 |
| **Mean with original MSAesm_ddG** | This study | **0.59** | 0.92 | 1.34 |
| **Fixed equal-weight ensemble** | This study | **0.59** | **0.91** | **1.33** |

We next evaluated PTmul-NR, containing 82 multiple-substitution variants on 13 distinct WT sequences, using the background-aware EvoPhi-DDG extension introduced above, which preserves whole-variant antisymmetry (Methods, Section 3.4). The EvoPhi-DDG extension achieved Pearson 0.66, compared with 0.61 for native JanusDDG. Their equal-weight mean reached 0.66 (Fig. 5j–l; Supplementary Table S9). The ensemble therefore improved on JanusDDG, but did not outperform the EvoPhi-DDG extension alone.

With only 13 WT sequences, these comparisons remain sensitive to the available protein sample.

Published PTmul-NR reference values include Pearson 0.59 for DDGemb, 0.36 for FoldX, 0.28 for MAESTRO, 0.23 for DDGun and 0.17 for DDGun3D [2]. These contextual values reinforce the relevance of the observed EvoPhi-DDG extension and ensemble, while not substituting for a matched rerun of all competing methods.

### 2.4 Classification of stabilizing and destabilizing variants

We next assessed whether the ensemble’s regression gains translated into improved discrimination between stabilizing and destabilizing variants, considering both ranking performance and classification at a fixed decision threshold. Following the criterion used for the JanusDDG stability-classification analysis on S461 [11], we excluded variants with experimental |ΔΔ*G*| ≤ 0.50 kcal/mol and retained only stabilizing (ΔΔ*G >* 0.50) or destabilizing (ΔΔ*G <* −0.50) variants. Here we apply this criterion to S669, S461, S96 and PTmul-NR. Variants are classified as stabilizing when predicted ΔΔ*G >* 0. To avoid threshold optimization or data selection bias, all predictions, including near-zero values, are retained in the evaluation (Methods, Section 3.6).

We compared JanusDDG, original MSAesm_ddG, EvoPhi-DDG and their antisymmetric equal-weight ensemble (Supplementary Table S10). On S669, the ensemble reached ROC-AUC 0.83, compared with 0.79 for JanusDDG, 0.82 for original MSAesm_ddG and 0.81 for EvoPhi-DDG (Fig. 6a,b). However, EvoPhi-DDG achieved higher recall and balanced accuracy than the ensemble, while original MSAesm_ddG had the highest precision. Averaging therefore improved ranking discrimination without uniformly improving classification at the fixed zero threshold.

**Figure 6:**
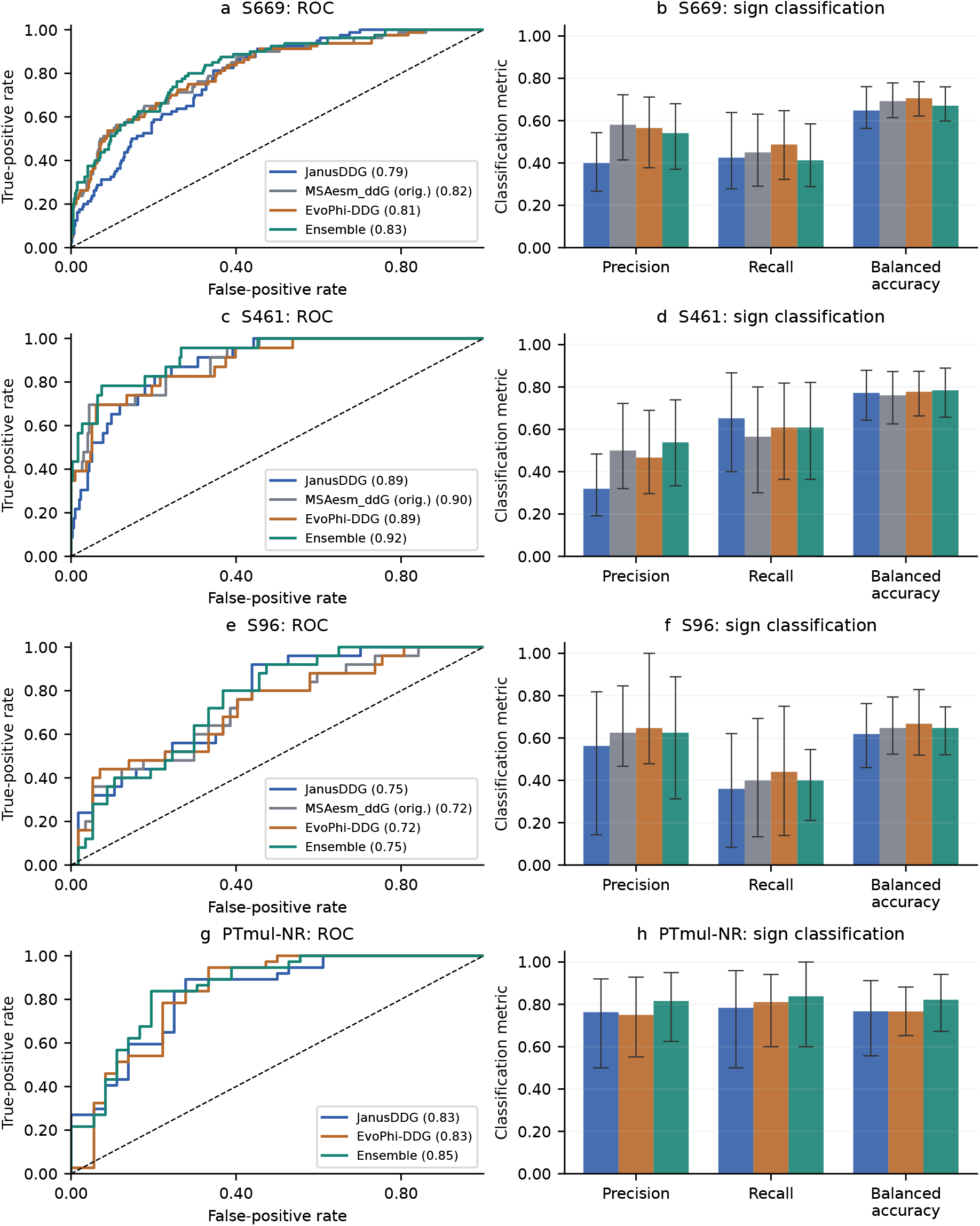
Stabilizing-versus-destabilizing classification for individual models and their ensemble. S669 (**a,b**), S461 (**c,d**), S96 (**e,f**) and PTmul-NR (**g,h**) after retaining only experimental ΔΔ*G >* 0.50 or ΔΔ*G <* −0.50 kcal/mol. ROC curves use continuous scores; legend parentheses show ROC-AUC. Bars show precision, recall and balanced accuracy at prediction threshold zero, with 95% percentile protein-bootstrap intervals from 1,000 paired draws. There are 81 retained WT sequence groups on S669, 44 on S461, 13 on S96 and 12 on PTmul-NR. Original MSAesm_ddG is included for the single-mutant comparison; the multiple-mutant EvoPhi-DDG score uses the background-aware construction preserving whole-variant antisymmetry.

On S461, the ensemble achieved ROC-AUC 0.92, compared with 0.89 for JanusDDG, 0.90 for original MSAesm_ddG and 0.89 for EvoPhi-DDG (Fig. 6c,d). Precision and balanced accuracy also improved relative to both components, while recall remained slightly lower than JanusDDG’s. Only 23 stabilizing variants were available, making these comparisons sensitive to individual examples.

On S96, ensemble ROC-AUC was 0.75, nearly unchanged from JanusDDG’s 0.75, although precision, recall and balanced accuracy increased. EvoPhi-DDG retained higher recall and balanced accuracy than the ensemble (Fig. 6e,f). Thus, the regression improvement did not translate into a substantial ranking improvement over JanusDDG or uniformly better classification than EvoPhi-DDG.

On PTmul-NR, the ensemble achieved precision 0.82, recall 0.84, balanced accuracy 0.82 and ROC-AUC 0.85, improving these point estimates relative to both JanusDDG and EvoPhi-DDG (Fig. 6g,h). The lack of a full-set regression advantage over EvoPhi-DDG did not preclude a benefit for this filtered classification task.

### 2.5 Accuracy through averaging, thermodynamic consistency through EvoPhi-DDG

To disentangle the effects of prediction averaging, antisymmetrization and hidden-bias removal, we compared the fixed mean of published antisymmetric JanusDDG and original directional MSAesm_ddG:

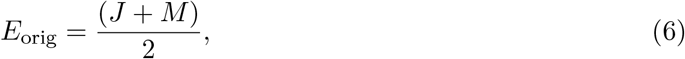

with the ensemble using antisymmetrized MSAesm_ddG with its published hidden biases:

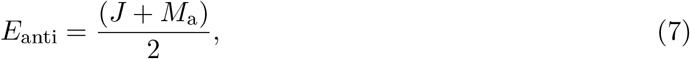

and the primary ensemble using EvoPhi-DDG:

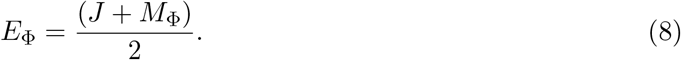

Published checkpoints, sequences, alignments, experimental labels and averaging weights were identical; EvoPhi-DDG additionally sets the hidden biases of the antisymmetric MSAesm_ddG head to zero (Methods, Section 3.3).

On S669, Pearson was identical at the reported precision: 0.59 for *E*_orig_, *E*_anti_ and *E*_Φ_. Thus the increase from JanusDDG Pearson 0.55 to approximately 0.59 is already obtained by averaging with the original MSAesm_ddG model; it cannot be attributed to antisymmetrization or hiddenbias removal. The same pattern was observed on S461 and S96, where the ensemble gains likewise arose from averaging rather than the potential-based reformulation (Supplementary Table S5).

Original-head ensemble ROC-AUC point estimates were slightly higher on all four filtered classification sets, while fixed-zero-threshold classification changes were metric dependent. We therefore do not interpret the potential-based reformulation as a general accuracy or ranking improvement.

The physical distinction was much larger. Original-head ensemble reversal MAEs were 0.10, 0.10 and 0.12 kcal/mol on S669, S461 and S96, respectively; both *E*_anti_ and *E*_Φ_ eliminated these defects by construction. On the identical S669 same-site pathways with nine intermediates, *E*_orig_ had mean direct-versus-path correlation 0.93 and closure RMSE 0.35 kcal/mol. With the antisymmetrized MSAesm_ddG, these became 0.98 and 0.17 kcal/mol, a 50.18% reduction in closure RMSE (Supplementary Fig. S2). Replacing this component with EvoPhi-DDG retained the displayed correlation and closure RMSE, while making the component’s same-site closure error exactly zero. Consequently, the signed closure error of *E*_Φ_ is exactly half that of JanusDDG under fixed mutation-site and alignment contexts. Original MSAesm_ddG averaging even had slightly lower pathway correlation than JanusDDG alone (0.94), despite its higher experimental Pearson. The controlled comparison therefore separates predictive complementarity from physical consistency.

For PTmul-NR, we compared four ways of constructing the MSA-derived prediction for a complete multiple-substitution variant, each combined with the same native JanusDDG prediction by equal-weight averaging (Supplementary Table S5). Let *A* be the complete WT sequence and *B* the fully mutated sequence. Within each endpoint-based sum, the starting background remains fixed: the forward sum scores each substitution independently from *A*, whereas the reverse sum scores each reversion independently from *B* (Methods, Section 3.4).

The *directional WT star* uses only the forward sum of original, directional MSAesm_ddG scores. Its ensemble reached Pearson 0.66 but retained a reversal MAE of 0.28 kcal/mol. The *whole-projected raw star* uses the same original single-substitution predictor, additionally computes the reverse sum on *B*, and takes half the difference between the two sums. This projects the complete MSA-derived variant score onto an antisymmetric form before combining it with JanusDDG; the individual single-substitution scores remain unprojected. The *component/whole projection* additionally antisymmetrizes every constituent single-substitution score by evaluating that transition in both directions, while retaining the published hidden biases. It then applies the same whole-variant projection to the forward and reverse endpoint sums. Finally, *JanusDDG + EvoPhi-DDG* uses this same component-and-whole construction after setting the MSAesm_ddG hidden biases to zero.

All four ensembles reached Pearson 0.66 at the reported precision; the three whole-projected constructions guaranteed exact whole-variant antisymmetry. Comparing the second and third rows isolates the effect of single-substitution projection at fixed endpoint backgrounds, and comparing the third and fourth isolates hidden-bias removal.

### 2.6 Residual complementarity explains when averaging helps

We examined the relationship between component errors rather than attributing ensemble gains solely to model diversity. On S669, JanusDDG and EvoPhi-DDG residuals were strongly correlated (*r* = 0.85), but had opposite signs for 24.96% of variants. The ensemble reduced absolute error below both individual predictors for 14.35% of variants (Supplementary Fig. S3). Averaging can also improve overall performance without being the best prediction for each individual case.

Writing *e*_*J*_ = *J* − *y* and *e*_Φ_ = *M*_Φ_ − *y*, the ensemble MSE decomposes exactly:

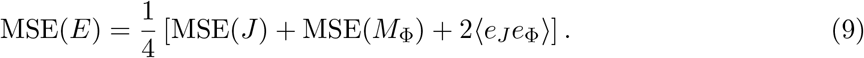

On S669 the component MSEs were 1.92 and 1.93, and the residual cross-product was 1.64, yielding ensemble MSE 1.78 in squared kcal/mol units. This quantifies the reduction made possible by imperfectly shared errors, despite their high correlation.

The curated S461 set showed lower component residual correlation (*r* = 0.62), opposite-sign errors for 29.28% of variants and an ensemble absolute error below both components for 16.70%. Its component MSEs were 0.92 and 0.94, with cross-product 0.61, yielding ensemble MSE 0.77. These descriptive differences indicate stronger error cancellation within the curated subset.

S96 component residuals had correlation 0.86; errors had opposite signs for 21.88% of variants, and the ensemble improved absolute error over both components for 9.38%. Its MSE decreased to 4.17, compared with 4.39 and 4.58 for JanusDDG and EvoPhi-DDG, respectively. The reduction is consistent with imperfectly shared errors, while the wide protein-bootstrap intervals limit conclusions about generalization.

On PTmul-NR, residual correlation was higher (*r* = 0.91), only 13.41% of variants had opposite-sign errors, and 7.32% improved over both components. Moreover, the EvoPhi-DDG extension was already more accurate than JanusDDG. These observations explain why a fixed mean improves JanusDDG but does not improve the stronger component on the full multiplemutant set. Supplementary Table S6 reports paired uncertainty for both comparisons on all four evaluated sets.

## 3 Methods

### 3.1 Datasets and multiple sequence alignment construction

#### 3.1.1 Datasets

The published JanusDDG and MSAesm_ddG checkpoints were developed using different training datasets. We retained the published weights without additional training, with the prescribed removal of the MSAesm_ddG hidden biases to obtain EvoPhi-DDG, and evaluated their predictions on common experimental records.

JanusDDG was trained on S2450 during both its initial supervised training and its subsequent transitivity-oriented fine-tuning [11]. S2450 was derived from S2648 by comparing full-length UniProt sequences and removing training proteins with greater than 25% sequence identity to any S669 protein [2]. This filtering removed 198 substitutions across 18 proteins, leaving 2,450 experimentally characterized single substitutions. The training data were augmented with the corresponding inverse substitutions using antisymmetry. The multiple-substitution dataset M28 was used to select the fine-tuning duration, rather than to optimize model parameters [11].

The original MSAesm_ddG checkpoint was trained on S155329, a filtered subset of the megascale stability dataset [10, 13]. This set contains 155,329 direct single substitutions across 149 protein backgrounds, following sequence-similarity filtering and exclusion of proteins without identified homologs in UniClust30. The authors performed local pairwise BLASTp alignments and excluded a training protein if an alignment with a test sequence simultaneously satisfied three conditions: sequence identity greater than 25%, E-value below 0.01, and alignment overlap greater than 50%. Identity was calculated relative to alignment length, while overlap was defined as alignment length divided by query length [10].

We evaluated three single-substitution benchmarks, S669, S461 and S96, and one multiple-substitution benchmark, PTmul-NR. We verified that all four datasets satisfy the sequence-similarity exclusion criterion used in the original MSAesm_ddG study: no evaluation sequence produced a BLASTp alignment against any of the 149 released training backgrounds simultaneously exhibiting sequence identity greater than 25%, query coverage greater than 50%, and an E-value below 0.01. Sequence mappings and experimental values were taken from the datasets distributed with JanusDDG to ensure comparisons on common evaluation records [11]. The individual benchmarks are described below.

S669 was introduced as a manually curated single-substitution benchmark with reduced sequence similarity to commonly used stability-training datasets [1]. We evaluate its 669 direct variants. Inverse variants are used for physical-consistency checks rather than pooled into the primary predictive-accuracy scores. In the full-length sequence mapping used here, the records correspond to 87 distinct WT sequences, which define the bootstrap resampling groups. Different protein identifiers mapping to the same WT sequence are therefore not treated as independent groups.

S461 is the published curated derivative of S669 [14]. We use the 461 direct-variant records, experimental values and full-length sequence mappings distributed with JanusDDG [11]. Its 48 protein identifiers correspond to 46 distinct WT sequences, used as resampling groups. Because S461 derives from S669, it provides a sensitivity analysis to dataset curation rather than an independent benchmark.

S96 contains 96 single-site variants across 14 distinct WT sequences, derived from ProTherm and manually checked against the original experimental reports [15].

PTmul-NR is a reduced multiple-substitution benchmark derived from PTmul [2, 6]. The distributed sequence table used here contains 82 variants across 13 distinct WT sequences, with two to five substitutions per variant.

#### 3.1.2 MSA generation and fixed alignment contexts

Multiple sequence alignments were generated using the public ColabFold MMseqs2 service [16, 17] and subsequently filtered with HHfilter [18]. Filtering used a maximum pairwise sequence identity of 90%, a diversity setting of 1000, minimum coverage and minimum query identity of 0, and a query-score threshold of −20.

To control memory requirements, alignment depth was restricted to a budget of 16,000 tokens. For an effective query length *L*, the number of retained rows, including the query, was

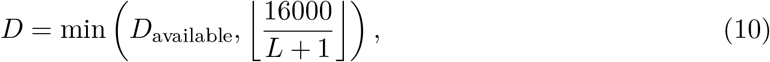

where *D*_available_ is the alignment depth after filtering and the additional token accounts for the initial sequence token. Rows were retained in their post-filtering order.

For S669 sequences exceeding the MSA input-length limit, we used a mutation-centered window containing at most 1,022 residues. Window selection depended only on sequence length and mutation position, without using experimental labels. This restriction affected 19 S669 variants. JanusDDG instead received the full sequences, which reached 3,685 residues in this dataset. S461 followed the same windowing and token-budget rules, reusing the verified alignment contexts for matching S669 variants. S96 and PTmul-NR sequences were within the MSA length limit and required no windowing. The same filtering procedure and token budget were applied across datasets.

To ensure consistent representations in forward–reverse and pathway comparisons, we held the alignment context fixed across the sequence states involved in each comparison. Specifically, homologous sequence rows, their order, alignment depth and window coordinates were unchanged; only the query row was modified to encode the substitutions present in the state being evaluated. Intermediate mutant states therefore reused the corresponding reference alignment rather than triggering new homolog searches or independent filtering. Within a fixed mutation site and alignment context, each sequence state consequently had the same representation wherever it appeared along a pathway, allowing prediction consistency to be assessed without introducing changes in the alignment context.

### 3.2 Models

This section describes the models used in this study, with particular emphasis on our potential-based EvoPhi-DDG reformulation of MSAesm_ddG and its extension to multiple substitutions while preserving whole-variant antisymmetry. We summarize the original JanusDDG and MSAesm_ddG architectures to provide the context needed for these developments; comprehensive architectural and training details are available in the corresponding publications [10, 11].

#### 3.2.1 MSAesm_ddG architecture

MSAesm_ddG predicts mutation-induced stability changes from contextual representations generated by MSA Transformer [9, 10]. Its input comprises the protein sequence of interest, referred to as the query sequence, together with an alignment of homologous sequences. The term query therefore identifies the sequence being evaluated within the alignment. The model extracts the query-sequence representations from the twelfth Transformer layer, which assigns a 768-dimensional embedding to each residue. These embeddings depend on both the query sequence and its alignment context.

Let *A* and *B* denote two protein sequences differing by a substitution at position *p*, and let C denote the fixed alignment context used to evaluate both sequences. We write

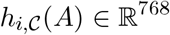

for the embedding of residue *i* in query sequence *A*. In particular, *h*_*p*,C_(*A*) represents the mutation site in the sequence and alignment context of *A*. A second vector summarizes the query sequence by averaging its residue embeddings:

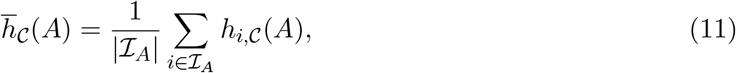

where I_*A*_ denotes the set of query positions included in the mean-pooling operation, following the original MSAesm_ddG implementation.

The mutation-site embedding and the mean-pooled query embedding are concatenated to form

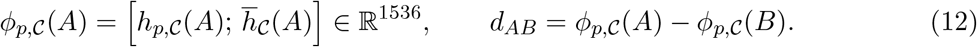

Here, the semicolon denotes concatenation of two 768-dimensional vectors. The representation therefore combines information at the mutation site with a summary of the full query sequence. Both vectors are recomputed for each endpoint sequence: fixing C holds the alignment context constant, but does not hold the query embeddings constant. The difference vector *d*_*AB*_ encodes the ordered comparison between *A* and *B*; reversing the endpoints gives *d*_*BA*_ = −*d*_*AB*_.

The prediction head maps this 1536-dimensional difference vector to a scalar through a single hidden layer containing 768 neurons:

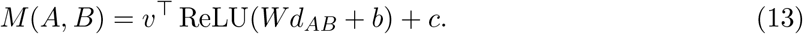

The matrix *W* ∈ ℝ^768×1536^ maps the input difference to the hidden layer, and *b* ∈ ℝ^768^ supplies one bias per hidden neuron. The vector *v* ∈ ℝ^768^ then combines the hidden activations into a scalar, to which the output bias *c* ∈ ℝ is added. All parameters are taken from the published checkpoint, and dropout is disabled during inference.

Although reversing the endpoints negates the input difference, the nonlinear prediction head does not generally satisfy *M* (*B, A*) = −*M* (*A, B*). Thus, the difference-based input representation alone does not guarantee antisymmetry of the predicted stability changes.

#### 3.2.2 JanusDDG architecture

JanusDDG is a physics-informed, sequence-based predictor that combines ESM-2 representations with a bidirectional cross-attention architecture [7, 11]. For two sequences *A* and *B* of length *L*, representing the wild-type and mutant proteins, respectively, the frozen ESM-2 encoder produces residue-embedding matrices

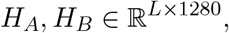

where each row contains the contextual representation of one residue. The directional backbone receives both the reference-state embedding *H*_*A*_ and the difference *H*_*A*_−*H*_*B*_, allowing it to process mutation-induced changes together with the sequence context in which they occur.

Two separate one-dimensional convolutions transform these inputs into 128-dimensional feature streams. Sinusoidal positional encodings retain information about sequence order. The streams then interact through two multi-head cross-attention operations: one uses the reference stream as queries and the difference stream as keys and values, while the other uses the difference stream as queries and the reference stream as keys and values. Queries and keys determine attention weights, which specify how the value vectors are combined. This mechanism couples the representation of the mutation-induced perturbation to that of the reference protein.

Residual connections, normalization and a position-wise feed-forward network integrate the attention outputs into 256-dimensional features at each sequence position. Global average pooling summarizes the mean activation of each feature across positions, whereas global maximum pooling retains its largest activation. Concatenating these summaries yields a 512-dimensional vector, which a scalar regression head maps to the directional prediction *F* (*A, B*).

Exact antisymmetry is imposed by evaluating the same backbone with shared parameters in both orientations:

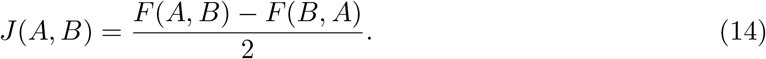

Thus, exchanging the endpoint sequences guarantees *J*(*B, A*) = −*J*(*A, B*), independently of the learned backbone parameters.

The published training procedure additionally promotes transitivity through a fine-tuning stage that combines supervised prediction with a consistency penalty:

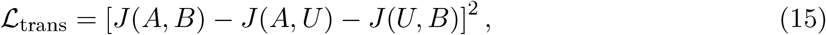

where *U* denotes a null reference represented by a zero embedding rather than a physical protein sequence. The notation *J*(*A, U*) denotes evaluation with this artificial embedding reference.

This penalty encourages the direct prediction to agree with the sum of predictions through a common reference, thereby promoting a state-potential interpretation of the predicted stability differences. Unlike antisymmetry, however, transitivity is encouraged through optimization rather than guaranteed algebraically for arbitrary intermediate sequences.

In this study, we use the published final checkpoint, retaining both its architectural antisymmetry and its transitivity-oriented fine-tuning.

#### 3.2.3 Antisymmetric siamese formulation and thermodynamic inconsistencies in MSAesm_ddG

A directional predictor *f* (*A, B*) estimates the stability change associated with the transition from protein sequence *A* to protein sequence *B*. Thermodynamic consistency requires the reverse transition to have the opposite predicted effect, *f* (*B, A*) = −*f* (*A, B*), and the identity transition to have zero effect, *f* (*A, A*) = 0. Neither condition is guaranteed for an arbitrary regression model.

To enforce these properties, JanusDDG applies the antisymmetric projection:

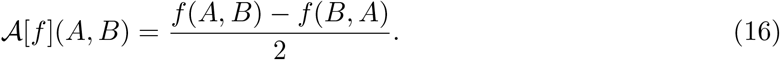

This operation combines the forward prediction with the sign-reversed backward prediction. Exchanging the endpoints therefore changes only the sign of the result:

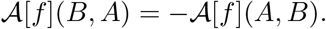

For identical endpoints, the two terms cancel, giving A[*f*](*A, A*) = 0. Equivalently, the projection removes the symmetric component [*f* (*A, B*) + *f* (*B, A*)]*/*2 and retains the antisymmetric component. It leaves an already antisymmetric predictor unchanged. The transformation is applied at inference using the existing model parameters; it requires neither additional training nor a newly fitted regression head.

Motivated by the simplicity and effectiveness of the antisymmetric projection used in Janus-DDG, we applied the same procedure to MSAesm_ddG, which does not enforce antisymmetry in its original formulation. To identify why this correction is needed for the original MSAesm_ddG predictor, consider its directional head,

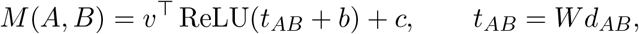

where *t*_*AB*_ is the linear transformation of the endpoint-representation difference before addition of the hidden bias and application of ReLU. Because *d*_*BA*_ = −*d*_*AB*_, reversing the transition gives *t*_*BA*_ = −*t*_*AB*_. However, ReLU is not an odd function: in general, ReLU(−*x*)/= − ReLU(*x*). Consequently, negating the input difference does not necessarily negate the prediction.

We quantify this violation by the reversal defect, defined as the sum of the forward and reverse predictions:

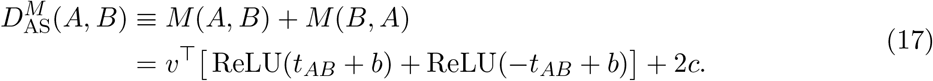

An exactly antisymmetric predictor has *D*^*M*^ (*A, B*) = 0 for every endpoint pair. The original head does not impose this condition. Likewise, identical endpoints give *d*_*AA*_ = 0 and hence

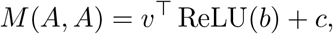

which is not constrained to vanish.

These violations are not attributable solely to the bias parameters. Even when *b* = 0 and *c* = 0, the identity ReLU(*x*) + ReLU(−*x*) = |*x*|, applied elementwise, gives

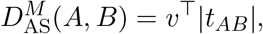

which is generally nonzero. Thus, removing the biases alone would not make the original directional head antisymmetric.

We next consider transitivity for three sequences *A, B* and *C*, evaluated at the same mutation position *p* and with the same alignment context C. Under these conditions, each sequence has a consistently defined representation *ϕ*_*p*,C_, so the endpoint differences satisfy

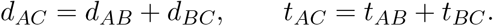

The input differences are therefore exactly additive. To determine whether this additivity is preserved by the prediction head, we define the triangle defect as

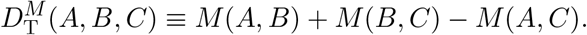

This quantity measures the discrepancy between the prediction along the two-step path *A* → *B* → *C* and the direct prediction for *A* → *C*. Substitution of the head expression yields

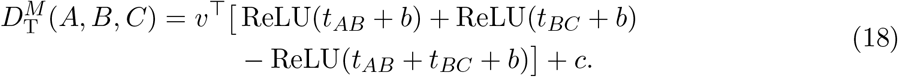

The remaining output-bias term is *c* because the two path predictions contribute 2*c*, whereas the direct prediction contributes *c*.

Even with zero biases, ReLU does not generally preserve additivity. Using ReLU(*x*) = (*x* + |*x*|)*/*2, the triangle defect becomes

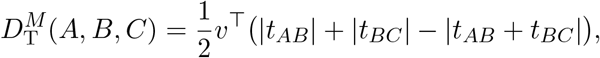

where absolute values are taken elementwise. This expression need not vanish, particularly when successive transitions have opposite signs in individual hidden coordinates. Thus, although end-point differencing guarantees additive input representations at fixed (*p*, C), the original nonlinear head does not guarantee either antisymmetry or transitivity. The antisymmetric projection enforces the former exactly; its consequences for transitivity require the separate analysis below.

#### 3.2.4 The antisymmetric MSAesm_ddG head is nearly transitive

The projected head is

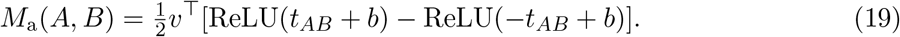

The scalar output bias cancels. If *b* = 0, the identity ReLU(*t*) − ReLU(−*t*) = *t* implies

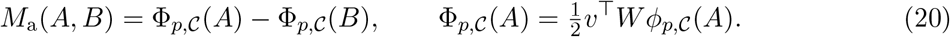

Hence all intermediate potentials telescope. This result requires a common state representation across edges, not a linear encoder.

For arbitrary hidden bias, ReLU(*x*) = (*x* + |*x*|)*/*2 yields the exact expression

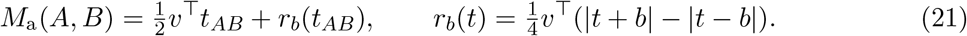

The residual is odd, preserving antisymmetry, but need not be additive. Consequently,

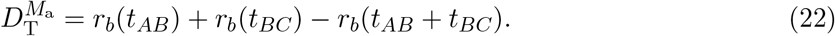

The reverse triangle inequality gives

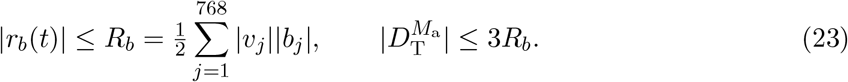

This is a sum of products of each neuron’s bias and output weight; it is not a sum of two scalar layer biases. For *N* intermediate states, the conservative closure bound is (*N* + 2)*R*_*b*_. In the checkpoint, *R*_*b*_ = 1.34 × 10^−2^ kcal/mol. These uniform bounds are looser than typical errors because edge residuals can cancel.

The proof applies to fixed position and alignment context. Changing the queried position, alignment homologs or window can change the representation of an intermediate state, preventing cancellation. The observed near-exact fixed-site transitivity therefore does not establish general multisite transitivity. Multisite paths are evaluated separately in Section 3.5.2, without extending that proof.

##### EvoPhi-DDG: exact same-site potential differences

We define EvoPhi-DDG by setting the hidden bias *b* to zero before applying the antisymmetric projection. The MSA Transformer, *W* and *v* retain their published weights; the scalar output bias *c* cancels under projection. Thus,

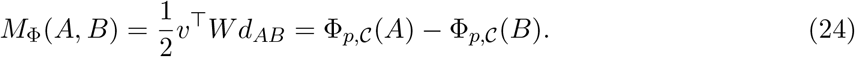

For any path with a common mutation position and alignment context, intermediate potentials cancel exactly, giving zero algebraic closure error. The original directional head and its anti-symmetric projection with the published nonzero hidden biases are retained as controls. This fixed-site guarantee does not imply general multisite transitivity.

### 3.3 Fixed ensemble and multiple-substitution construction

To integrate the physics-informed predictions of JanusDDG with the alignment-derived evolutionary information used by MSAesm_ddG, we combined the two predictors at the output level. This approach retains their independently learned representations and prediction functions, allowing their complementary information to contribute without training a joint architecture. For a single-substitution transition from sequence *A* to sequence *B*, we define

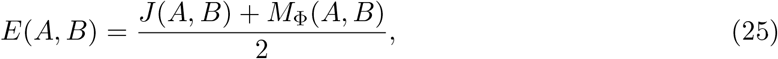

where *J* denotes the published JanusDDG predictor and *M*_Φ_ denotes EvoPhi-DDG (Equation 24). Both components predict the same quantity in kcal/mol and use the same sign convention. Equal weighting therefore preserves the prediction scale without introducing a fitted mixing coefficient or an additional calibration model.

The fixed average provides a deliberately parsimonious integration strategy in a setting where experimental stability measurements remain limited in their coverage of diverse, full-length proteins. Large-scale measurements on small domains have substantially expanded the available data, but their transfer to longer protein scaffolds remains challenging [19]. Learning a joint predictor or optimizing combination weights would introduce additional parameters and model-selection decisions supported by this limited experimental coverage. In contrast, after the prescribed hidden-bias removal, our construction leaves all remaining component parameters unchanged and requires no additional supervised fitting, avoiding the associated opportunity for overfitting at the combination stage.

This choice is also motivated by evidence that estimating combination weights can introduce sufficient sampling variability to offset their potential advantage over equal weighting [20]. Furthermore, studies of multimodal learning show that jointly trained information streams can exhibit different overfitting and generalization behaviors, complicating their effective integration [21]. These considerations motivate the fixed ensemble as a low-complexity strategy; they do not establish its superiority over a jointly trained model.

The construction also preserves the exact antisymmetry of its components. When both predictors are evaluated consistently under endpoint reversal,

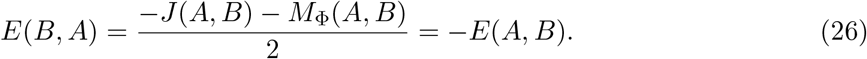

Thus, combining the predictions introduces no reversal inconsistency.

For the original-head control, define *E*_orig_ = (*J* + *M*)*/*2, and let *E*_anti_ = (*J* + *M*_a_)*/*2 use the published-hidden-bias antisymmetric MSAesm_ddG head, *M*_a_ = A[*M*]. Since JanusDDG is exactly antisymmetric,

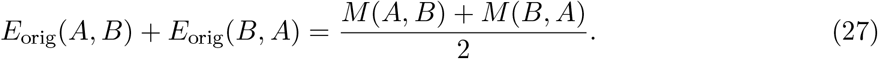

Thus, averaging with JanusDDG halves the original MSAesm_ddG reversal defect, whereas projecting the MSA head eliminates it. Linearity of the projection also gives

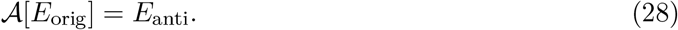

These controls retain the same JanusDDG predictions and equal averaging weights; the primary ensemble additionally replaces *M*_a_ with *M*_Φ_, isolating hidden-bias removal. Same-site physical-consistency comparisons reused the same S669 site/window contexts and the same random-residue paths across ten repetitions per intermediate count. JanusDDG edge predictions were held fixed while the original MSAesm_ddG head, its antisymmetric projection and EvoPhi-DDG supplied the alternative MSA contributions.

For a shared path *P* = (*S*_0_ = *A, S*_1_, … , *S*_*K*_ = *B*), define the signed closure error of a predictor *f* as

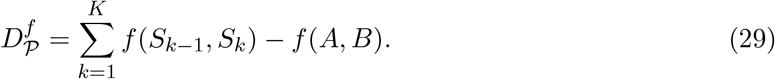

Linearity of the fixed average gives

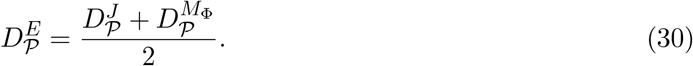

For fixed-site paths under a common alignment context, 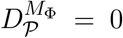, so the ensemble closure MAE and RMSE are exactly half those of JanusDDG in exact arithmetic. For multisite paths, a component with small closure error can reduce the pathway discrepancy of the other component, while oppositely signed closure errors can partially cancel.

### 3.4 Prediction of multiple-substitution effects

JanusDDG is designed to predict the effects of both single and multiple substitutions, whereas MSAesm_ddG was developed for single substitutions. To extend the ensemble to multiple-substitution variants, we therefore extended EvoPhi-DDG using an endpoint-based construction that preserves whole-variant antisymmetry. We also evaluated a sequential construction to assess sensitivity to the sequence backgrounds used to score the individual substitutions.

#### 3.4.1 Endpoint-based construction

Let *A* denote the complete WT sequence, *B* the complete multiple mutant, and *S* the set of substituted positions. For each *i* ∈ *S*, let *A*_*i*_ denote the sequence obtained by introducing only substitution *i* into *A*, and let *B*_−*i*_ denote the sequence obtained by reverting only substitution *i* in *B*. Thus, *B*_−*i*_ retains all substitutions except *i*.

Each single-substitution transition is scored using EvoPhi-DDG, *M*_Φ_, evaluated on the actual endpoint sequences of that transition. We compute

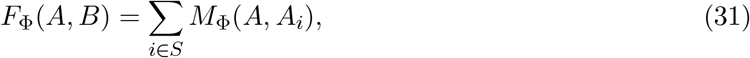

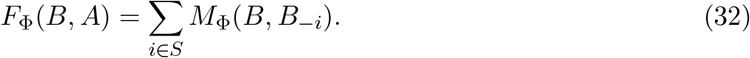

The first sum evaluates each substitution independently on the WT background, whereas the second evaluates each reversion independently on the fully mutated background. Neither sum represents a sequential path: the starting sequence remains fixed within each sum. The construction therefore considers each substitution both in the absence and in the presence of all other substitutions, without specifying a mutation order.

The whole-variant prediction is

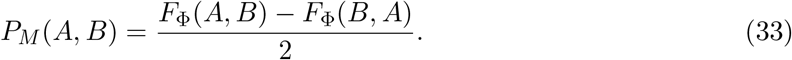

Exchanging the complete endpoint sequences interchanges the two sums, guaranteeing *P*_*M*_ (*B, A*) = −*P*_*M*_ (*A, B*). This property does not follow from antisymmetrizing individual WT-based contributions alone: under endpoint reversal, the reference background becomes the fully mutated sequence, and the corresponding contributions must be recomputed on that background.

The multiple-mutant ensemble combines *P*_*M*_ with the direct JanusDDG prediction between the complete endpoint sequences:

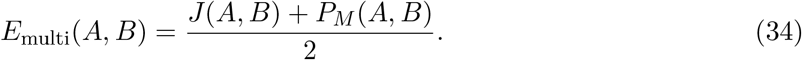

Both components are globally antisymmetric, so their fixed equal-weight average preserves this property.

#### 3.4.2 Sequential construction

As an alternative, we evaluated paths that update the sequence background after each substitution. For an ordering *π* = (*i*_1_, … , *i*_*k*_) of the *k* = |*S*| substituted positions, let *A*^(0)^ = *A* and obtain *A*^(*j*)^ by introducing substitution *i*_*j*_ into *A*^(*j*−1)^, such that *A*^(*k*)^ = *B*. The path prediction is

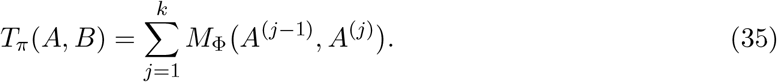

Unlike the endpoint-based construction, this procedure scores transitions on partially mutated backgrounds. Its result can depend on substitution order because consistency at a fixed mutation site does not guarantee path independence when successive transitions involve different sites.

We used the ordering *π*_↑_, defined by increasing sequence position, for both forward substitutions and reverse substitutions. The sequential whole-variant prediction is

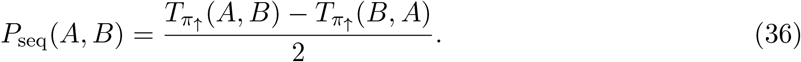

Applying substitutions in increasing position order in both directions does not generally traverse the same path in reverse. The final projection therefore ensures whole-variant antisymmetry independently of any residual pathway dependence. The order was fixed without using experimental test labels.

### 3.5 Thermodynamic-consistency evaluation

We assessed thermodynamic consistency separately from predictive accuracy by comparing direct predictions with sums along alternative mutation paths. We first considered paths confined to a single substitution site, then evaluated paths involving temporary substitutions at other sites. In both tests, the endpoint sequences differ by a single substitution. The multisite paths therefore assess pathway consistency independently of the construction used to predict effects between endpoints differing at multiple positions.

For a predictor *f* and a path *P* = (*X*_0_, … , *X*_*K*_), we define the signed closure error as

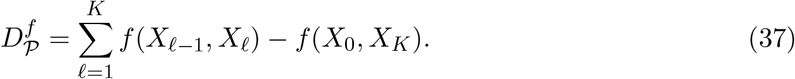

A zero closure error indicates agreement between the direct prediction and the sum along the alternative path. This agreement concerns internal prediction consistency and does not require experimental stability measurements for the intermediate states.

#### 3.5.1 Same-site pathways

We followed the random-residue pathway construction used to evaluate JanusDDG [11]. For each S669 variant, we retained the endpoint sequences and sampled *N* = 1, 3, 5, 7, 9 intermediate residues uniformly from the 20 standard amino acids, with replacement. Only the residue at the target mutation site was changed along each path; the remainder of the sequence was held fixed. Endpoint residues were eligible for sampling, and consecutive identical states were allowed.

For each value of *N* , ten repetitions were generated using a fixed seed, with identical paths shared across predictors. These newly generated paths are not assumed to reproduce the original authors’ unreleased random draws. For each repetition, we computed the Pearson correlation between direct and path-summed predictions across all 669 variants, together with closure MAE, RMSE and signed bias.

For the MSAesm_ddG calculations, all 20 residue states were encoded for each of the distinct site/window contexts. Each encoded state was reused identically wherever it occurred, ensuring that comparisons reflected the prediction head rather than changes in the representation of a shared state. The original head, its antisymmetric projection with the published hidden biases, and EvoPhi-DDG were evaluated on exactly the same encoded states. Reversal defects were assessed separately, including predictions for identity transitions.

#### 3.5.2 Multisite pathways on S669

For each of the 669 S669 WT–mutant pairs (*A, B*), differing at target position *p*, we sampled paths with *N* = 2, 4, 6, 8, 10 intermediate states. For *m* = *N/*2, we sampled *m* distinct non-target positions uniformly from positions occupied by standard amino acids within the existing MSA window. At each temporary site, the replacement was sampled uniformly from the 19 standard amino acids different from its WT residue. Temporary substitutions were introduced in the sampled order; the target substitution was then applied, followed by reversion of the temporary substitutions in reverse order. Thus, every path starts at *A*, ends at *B*, and contains *N* + 1 single-substitution edges. All temporary substitutions are absent from the final state.

For each intermediate count, ten repetitions were generated with a fixed seed of 42, and the same paths were shared across predictors. This yielded 6,690 paths per intermediate count and 33,450 paths overall. Sequence states were deduplicated within their reference MSA context, giving 196,196 context-specific states in 92 query/window contexts and 229,558 distinct directed edges. Temporary sites and replacement residues were sampled without using experimental stability measurements.

All query states within a reference MSA context shared homolog rows, row order, depth and window coordinates, following Section 3.1.2. Only the query sequence was changed. The original MSAesm_ddG head, its antisymmetric projection and EvoPhi-DDG used the same frozen encoded states. JanusDDG used frozen ESM-2 representations of the corresponding full-length sequence states, with temporary-site coordinates mapped from each MSA window to the full sequence. Reused sequence states had identical representations wherever they occurred. We verified JanusDDG edge evaluation against the unmodified published checkpoint and direct S669 predictions against the existing full-length prediction cache. No new homolog searches, training, score calibration or ensemble fitting were performed.

For each predictor *f* , the direct prediction *f* (*A, B*) was compared with the sum over the *N* + 1 path edges using the signed closure error defined above. Every edge is a single substitution. The ensemble therefore averages JanusDDG and EvoPhi-DDG on each edge, rather than invoking the multiple-substitution endpoint construction *E*_multi_.

This test probes a property not guaranteed by the fixed-site potential explanation. Partitioning the linear functional of the EvoPhi-DDG head into mutation-site and pooled contributions gives

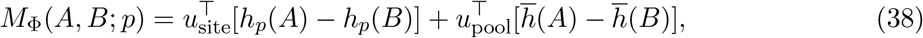

where *u*_site_ and *u*_pool_ are the two blocks of *W* ^⊤^*v/*2. The antisymmetric control retaining the published hidden biases additionally contains the bounded residual *r*_*b*_ described above. Under a fixed alignment context, the pooled contribution telescopes along a path. However, the site-specific term uses the embedding at the position changed by each transition. When that position changes between edges, the fixed-site argument no longer establishes a common global potential. Multisite closure is therefore evaluated empirically.

For each repetition and intermediate count, we computed Pearson correlation between direct and path-summed predictions across all 669 variants, together with closure MAE and RMSE. Reported means and standard deviations describe the ten repetitions; maximum absolute defects are taken over all 6,690 variant–repetition pairs at the corresponding intermediate count. Experimental labels were not used in this consistency analysis.

Descriptive uncertainty in ensemble-minus-component comparisons was estimated from 5,000 paired bootstrap resamples of the 87 complete WT-protein groups. All variants and all ten repetitions belonging to each sampled protein were retained together. Within each bootstrap draw, RMSE was calculated separately for each repetition and then averaged, matching the reported point estimate. Identical draws were used for all predictors. The within-protein comparison pooled squared closure errors across variants and repetitions before taking the square root.

### 3.6 Binary classification excluding the experimental uncertainty band

Following the label-filtering strategy of JanusDDG [11], define the common retained set *I* = {*i* : *y*_*i*_ *>* 0.50 or *y*_*i*_ *<* −0.50}, with *y*_*i*_ in kcal/mol. Values exactly equal to ±0.50 are excluded, matching the strict inequalities of the released classification procedure. The positive class is *z*_*i*_ = **1**[*y*_*i*_ *>* 0.50]. Every selected example is classified by 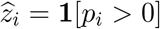; zero predictions map to the negative class. This separates the experimental exclusion band from the prediction decision boundary. We do not remove predictions with |*p*_*i*_| ≤ 0.50, and the ensemble and its components share identical retained examples.

For true-positive, false-positive, true-negative and false-negative counts TP, FP, TN and FN, the principal metrics are

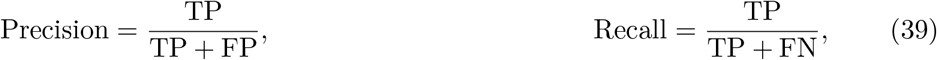

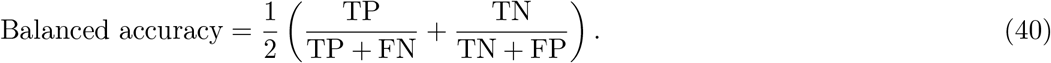

ROC-AUC is calculated from the continuous regression scores.

Classification intervals use 1,000 paired resamples of retained WT sequence groups with replacement. These are 81 groups for S669, 44 for S461, 13 for S96 and 12 for PTmul-NR. The same indices are used for every model and for ensemble-minus-component comparisons. Error bars are percentile protein-bootstrap intervals, rather than treating individual mutations as independent observations.

### 3.7 Predictive statistics

Predictive accuracy was assessed using Pearson and Spearman correlations, MAE and RMSE against experimental ΔΔ*G* values.

Descriptive uncertainty intervals were estimated using 5,000 paired bootstrap resamples of groups defined by identical complete WT sequences. Groups were sampled with replacement, retaining all associated variants each time a group was selected. Identical resamples were used for each ensemble and its comparator to preserve the pairing when estimating performance differences.

For ensemble-control comparisons, Pearson differences were defined as *r*_anti_ − *r*_orig_, whereas MAE and RMSE reductions were defined as the original-head ensemble error minus the corresponding antisymmetric-head ensemble error. Positive values therefore favor the antisymmetric-head ensemble for all three comparisons. On PTmul-NR, the comparison between the *whole-projected raw star* and *component/whole projection* constructions used the same paired protein-bootstrap procedure, holding both endpoint backgrounds and whole-variant projection fixed. EvoPhi-DDG ensemble differences relative to the original-head ensemble were evaluated analogously. Classification comparisons followed the same experimental inclusion rule |ΔΔ*G*| *>* 0.50 kcal/mol and fixed prediction threshold of zero described in Section 3.6.

## 4 Discussion

Our results show that an existing alignment-conditioned predictor can be reformulated as a site-conditioned stability potential without additional training and then integrated with the previously published JanusDDG model to improve stability predictions. A fixed, equal-weight average out-performed both components on the single-substitution benchmarks, supporting the presence of complementary predictive information despite substantial correlation between their errors. The potential-based reformulation provides a distinct benefit: it preserves predictive performance while enforcing exact reversal consistency and exact same-site transitivity under fixed alignment contexts. Exact transitivity is not guaranteed for arbitrary multisite paths or for the ensemble. Together, these findings identify complementary representations and physical constraints as two separable contributions to the resulting predictor.

The predictive gains suggest that neither component fully captures the information available to the other. On S669, the ensemble increased Pearson correlation by 0.05 over JanusDDG and reduced MAE and RMSE relative to both components. Residual analysis provides a statistical explanation: although component errors were strongly correlated, their differences allowed partial error cancellation. The gain persisted on S461, where Pearson correlation reached 0.75, and on S96, where it reached 0.55. These observations are consistent with a benefit from combining contextual single-sequence representations with explicit alignment-derived evolutionary information. However, the predictors also differ in architecture and training data, so these comparisons establish complementarity between their predictions rather than isolating the causal contribution of alignment information alone.

The fixed combination is particularly relevant given the limited experimental coverage of diverse, full-length proteins. It exploits existing predictors without estimating additional combination weights, calibrating their outputs or training a joint architecture. This avoids additional fitting at the integration stage, although it does not remove limitations inherited from the component models or establish superiority over a jointly trained alternative. The present results therefore demonstrate the value of a parsimonious integration strategy rather than the optimality of equal weighting.

The potential-based formulation has an explicit architectural explanation. Antisymmetry alone does not imply transitivity. For the specific single-hidden-layer ReLU head considered here, however, the projection yields an exact potential difference when the hidden biases vanish. Nonzero biases introduce a bounded nonlinear residual, explaining why the published head can exhibit nearly exact transitivity at a fixed mutation site and alignment context after projection. This connects the observed improvement to an identifiable property of the prediction head rather than to a general assumption about antisymmetric networks.

The original-head controls distinguish this physical contribution from the predictive benefit of averaging. Combining JanusDDG with the original MSAesm_ddG predictor already produced the S669 correlation gain. Antisymmetrization retained essentially the same accuracy while eliminating the ensemble reversal defect and approximately halving its nine-intermediate pathway-closure RMSE. Removing the hidden biases further eliminates the component’s residual same-site closure error, yielding the exact potential-based formulation of EvoPhi-DDG without materially changing predictive accuracy. Consequently, under fixed mutation-site and alignment contexts, the ensemble’s signed closure error is exactly half that of JanusDDG. Across S669, S461 and S96, changes in predictive accuracy were small and did not show a consistent direction. The projection is therefore supported primarily by its exact reversal guarantee and improved pathway consistency, rather than by a uniform increase in regression or classification performance.

The S669 multisite paths show that these consistency gains extend beyond paths confined to one position. The ensemble reduced mean closure RMSE relative to JanusDDG at all five tested path lengths. With ten intermediates, within-protein RMSE improved in 68 of 87 WT groups, and the descriptive paired bootstrap interval favored the ensemble over JanusDDG. Its mean RMSE was also lower than EvoPhi-DDG at four of five path lengths, although the ten-intermediate bootstrap interval for this comparison included zero. The component signed closure errors were weakly correlated at ten intermediates (*r* = 0.02), supporting partial cancellation under averaging. Nevertheless, the site-dependent readout of EvoPhi-DDG prevents the fixed-site potential argument from establishing a common global potential. Residual multisite defects therefore remain, and the results support improved consistency over the evaluated paths rather than exact path independence. Because all path edges and endpoint pairs involve single substitutions, this conclusion is separate from the construction used to predict complete multiple-substitution variants.

Predictive performance on multiple substitutions illustrates a complementary limitation of fixed averaging. The whole-variant-antisymmetric EvoPhi-DDG extension was already more accurate than JanusDDG on the full PTmul-NR set, and the ensemble improved JanusDDG without surpassing the stronger component. This behavior is consistent with both the greater correlation of component residuals and their unequal predictive performance.

Classification further distinguishes predictive ranking from decisions at a fixed threshold. Out-side the experimental near-neutral band, improved ROC-AUC did not consistently translate into greater sensitivity to stabilizing variants at the prespecified zero threshold. On S669, for example, the ensemble improved ranking discrimination while retaining lower stabilizing-variant recall than EvoPhi-DDG. On S461, its ROC-AUC reached 0.92, but only 23 stabilizing examples were available. Conversely, on PTmul-NR, the ensemble improved precision, recall, balanced accuracy and ROC-AUC point estimates relative to both components. These differences show that regression accuracy, ranking discrimination and thresholded classification capture distinct aspects of performance.

This study distinguishes the physical guarantees obtained from a potential-based reformulation from the accuracy gains obtained by combining complementary predictions. EvoPhi-DDG supplies an alignment-conditioned component with exact same-site consistency under fixed contexts and a whole-variant-antisymmetric extension to multiple substitutions. Its integration with the original JanusDDG predictor illustrates how existing representations can be combined without further training, while keeping predictive accuracy and thermodynamic consistency as separate evaluation criteria.

## Data and code availability

The source code for EvoPhi-DDG and its ensemble with JanusDDG is available at https://github.com/bardu98/EvoPhi-DDG. The repository includes instructions for reproducing all results reported in this study.

## Supplementary Figures

**Figure S1:**
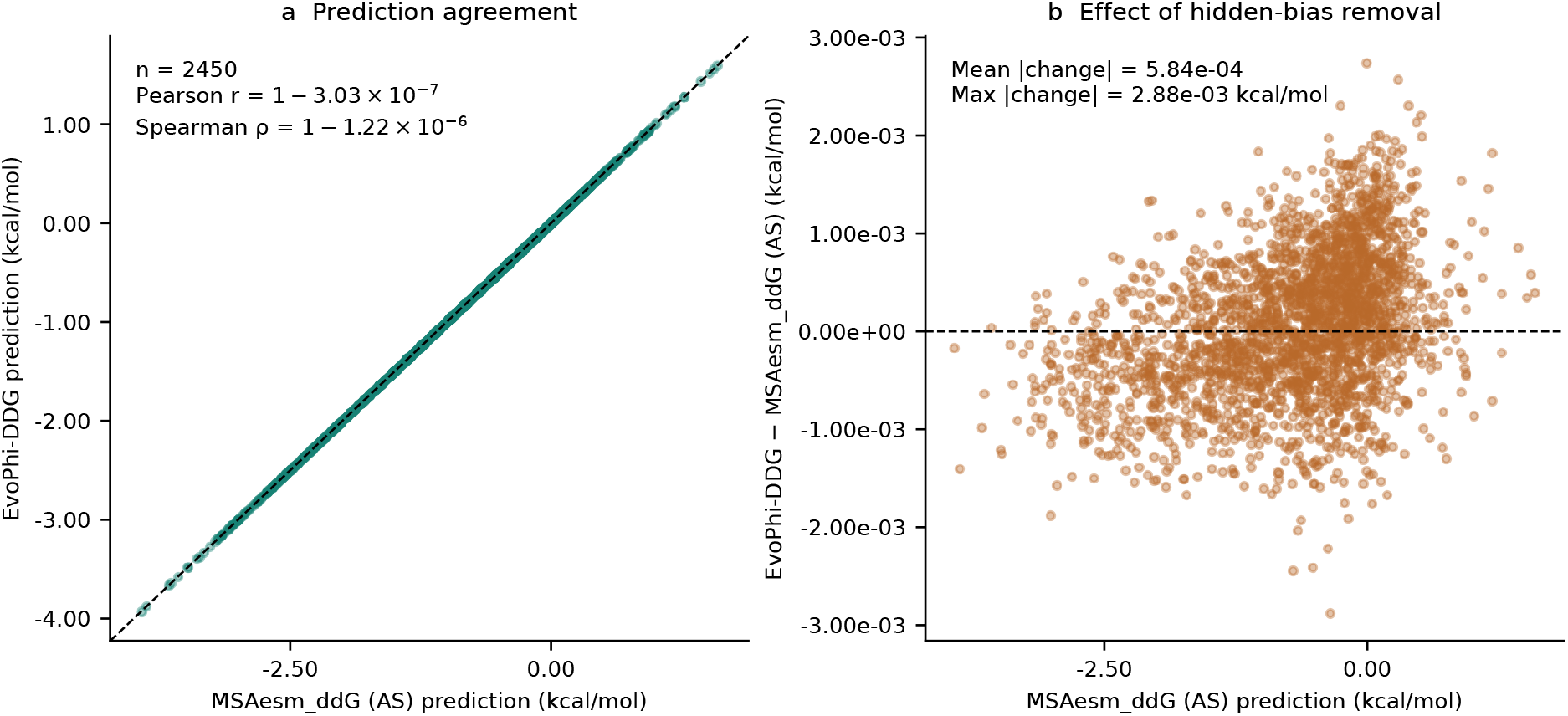
Effect of removing hidden biases on predicted stability changes. **a**, EvoPhi-DDG predictions versus those of antisymmetrized MSAesm_ddG with the published hidden biases, evaluated on S2450. The two formulations yielded Pearson and Spearman correlations of 1 − 3.03 × 10^−7^ and 1 − 1.22 × 10^−6^, respectively. **b**, Signed prediction differences (EvoPhi-DDG minus antisymmetrized MSAesm_ddG) as a function of the corresponding prediction with the published hidden biases. The mean and maximum absolute differences were 5.84 × 10^−4^ and 2.88 × 10^−3^ kcal mol^−1^, respectively.

**Figure S2:**
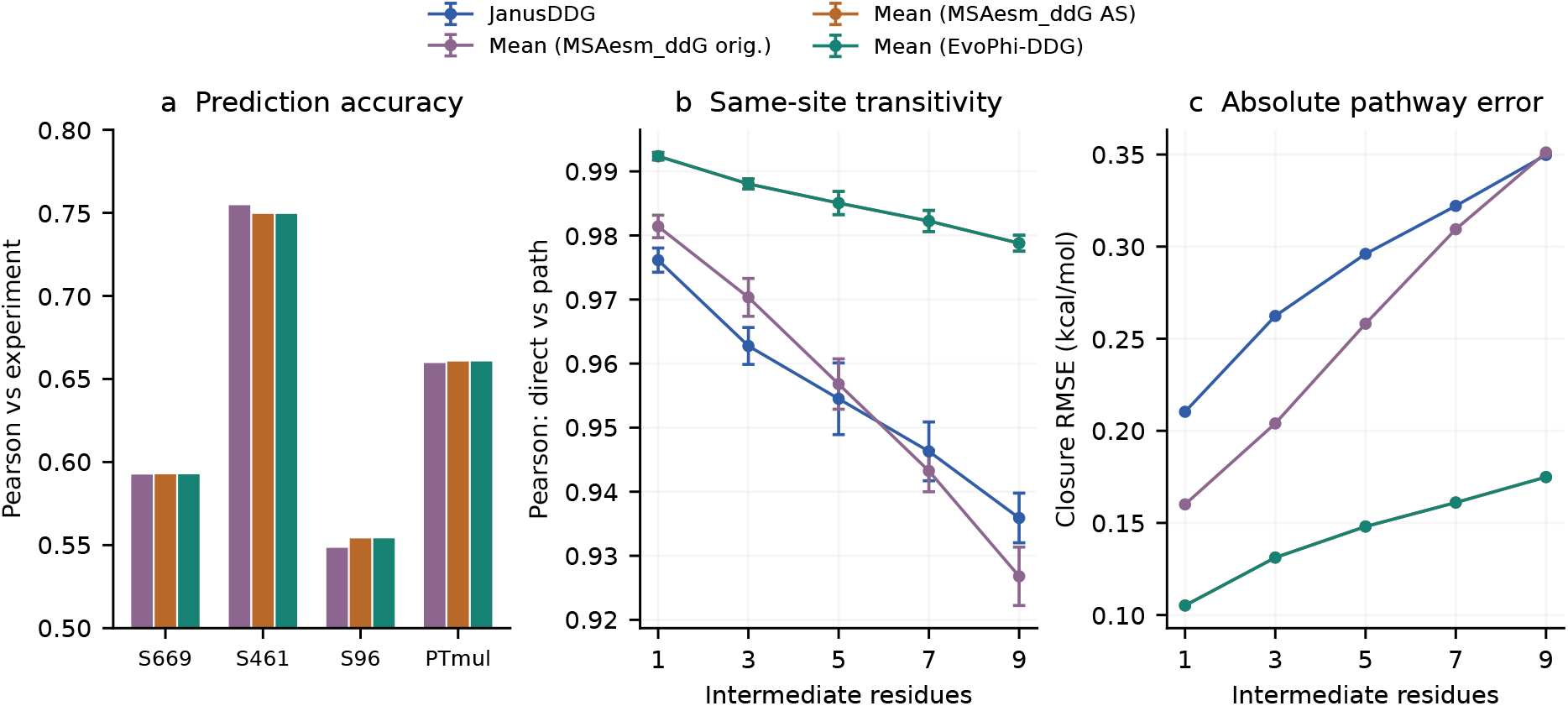
Distinguishing averaging from physical projection. **a**, Experimental Pearson of fixed original-MSAesm_ddG and antisymmetric ensembles. The PTmul-NR original baseline is a directional WT star sum; its projected counterpart additionally includes complete-mutant backgrounds. **b**, S669 same-site direct-versus-path correlation on identical states and paths; bars are standard deviations over ten repetitions. **c**, Closure RMSE on those paths. Accurate averaging alone does not guarantee reversal consistency or low pathway error.

**Figure S3:**
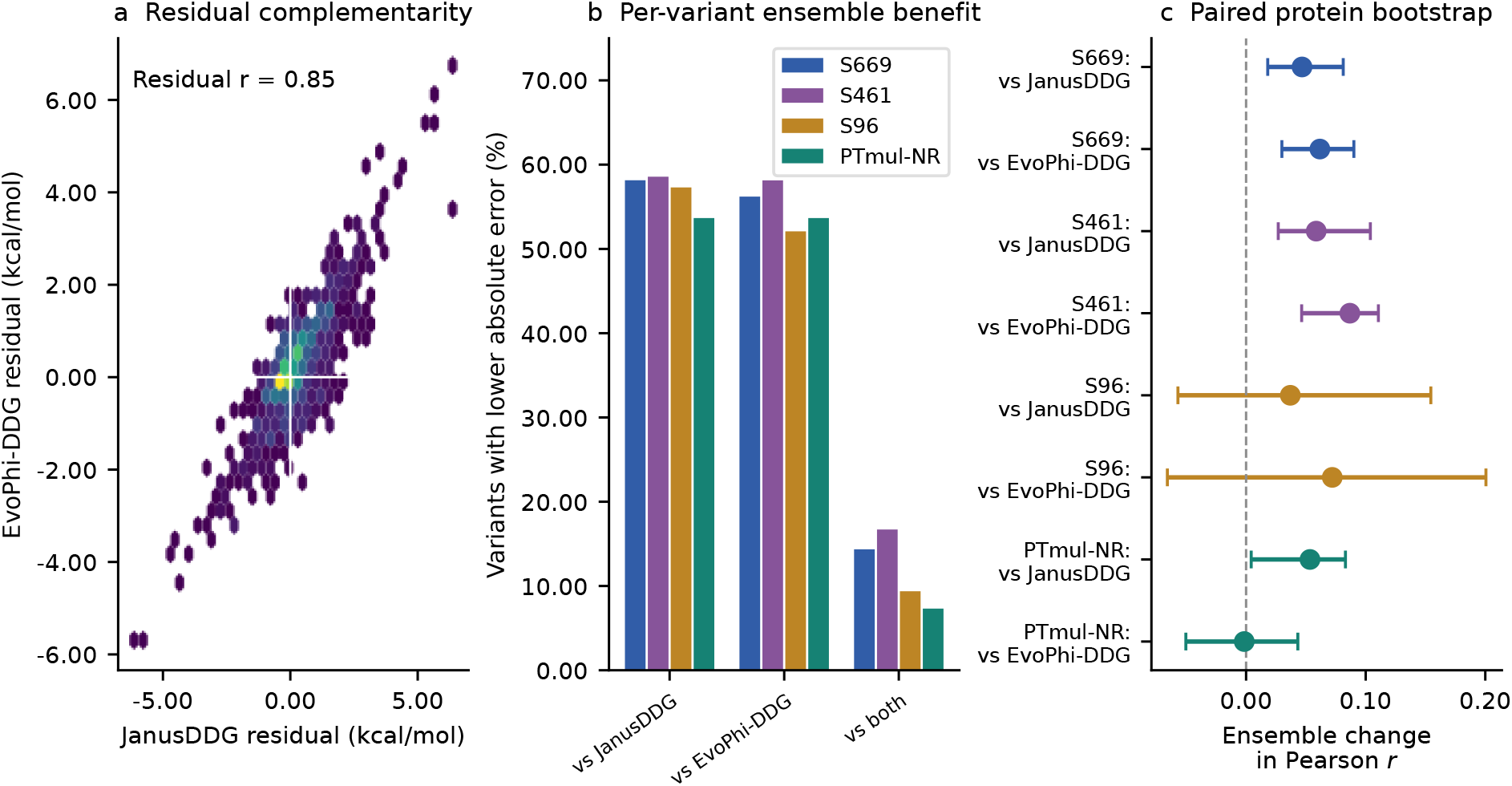
Complementarity and uncertainty of ensemble gains. **a**, S669 residuals of JanusDDG and EvoPhi-DDG; opposite-sign quadrants permit error cancellation. **b**, Percentage of variants where the ensemble reduces absolute error relative to Janus, EvoPhi-DDG, or both. The bars are overlapping comparisons, not disjoint categories. **c**, Ensemble-minus-component Pearson differences with paired 95% percentile protein-bootstrap intervals. Resampling uses 87 WT sequence groups for S669, 46 for S461, 14 for S96 and 13 for PTmul-NR, with 5,000 draws. The single-mutant sets overlap; their intervals are separate descriptive analyses.

## Supplementary Tables

**Table S1:** Detailed evaluation on S669. MAE and RMSE are in kcal/mol. Bold marks the best values and associated methods, including ties at two decimals.

| Predictor | Pearson | Spearman | MAE | RMSE |
|---|---|---|---|---|
| **S669** (*n* = 669) |  |  |  |  |
| JanusDDG | 0.55 | 0.55 | 0.97 | 1.39 |
| MSAesm_ddG original | 0.54 | 0.55 | 0.97 | 1.38 |
| MSAesm_ddG antisymmetrized | 0.53 | 0.55 | 0.99 | 1.39 |
| EvoPhi-DDG | 0.53 | 0.55 | 0.99 | 1.39 |
| **Equal-weight ensemble** | **0.59** | **0.61** | **0.91** | **1.33** |

**Table S2:** Detailed evaluation on S461. MAE and RMSE are in kcal/mol. Bold marks the best values and associated methods, including ties at two decimals.

| Predictor | Pearson | Spearman | MAE | RMSE |
|---|---|---|---|---|
| **S461** (*n* = 461) |  |  |  |  |
| JanusDDG | 0.69 | 0.66 | 0.71 | 0.96 |
| MSAesm_ddG original | 0.69 | 0.66 | 0.71 | 0.94 |
| MSAesm_ddG antisymmetrized | 0.66 | 0.65 | 0.73 | 0.97 |
| EvoPhi-DDG | 0.66 | 0.65 | 0.73 | 0.97 |
| **Equal-weight ensemble** | **0.75** | **0.72** | **0.65** | **0.88** |

**Table S3:** Detailed evaluation on S96. MAE and RMSE are in kcal/mol. Bold marks the best values and associated methods, including ties at two decimals.

| Predictor | Pearson | Spearman | MAE | RMSE |
|---|---|---|---|---|
| **S96** (*n* = 96) |  |  |  |  |
| JanusDDG | 0.52 | 0.50 | 1.50 | 2.10 |
| MSAesm_ddG original | 0.47 | 0.48 | 1.53 | 2.16 |
| MSAesm_ddG antisymmetrized | 0.48 | 0.47 | 1.53 | 2.14 |
| EvoPhi-DDG | 0.48 | 0.47 | 1.53 | 2.14 |
| **Equal-weight ensemble** | **0.55** | **0.52** | **1.45** | **2.04** |

**Table S4:** Detailed evaluation on PTmul-NR. MAE and RMSE are in kcal/mol. Bold marks the best values and associated methods, including ties at two decimals. The primary EvoPhi-DDG prediction uses zero hidden biases and component-and-whole antisymmetrization; the other constructions are sensitivity analyses.

| Predictor | Pearson | MAE | RMSE |
|---|---|---|---|
| JanusDDG | 0.61 | 1.57 | 2.06 |
| **MSAesm_ddG antisymmetrized** | **0.66** | **1.45** | **1.95** |
| **EvoPhi-DDG** | **0.66** | **1.45** | **1.95** |
| **Equal-weight ensemble** | **0.66** | 1.47 | **1.95** |
| **EvoPhi-DDG: sequential, both directions** | **0.66** | **1.45** | **1.95** |

**Table S5:** Fixed ensembles with original MSAesm_ddG, its antisymmetric projection and EvoPhi-DDG. MAE, RMSE and reversal MAE are in kcal/mol. Reversal MAE is ⟨|*E*(*A, B*)+*E*(*B, A*)|⟩. ROC-AUC uses the existing experimental |ΔΔ*G*| *>* 0.50 subsets: 474 S669, 319 S461, 82 S96 and 73 PTmul-NR variants. For single mutants, head projection and prescribed hidden-bias removal distinguish the controls. For PTmul-NR, every row is the equal-weight mean of the same native JanusDDG whole-variant prediction and one of four MSA-derived scores. With *A* denoting the complete WT and *B* the fully mutated sequence: (i) *directional WT star* sums original directional MSAesm_ddG predictions for substitutions introduced independently into *A*; (ii) *whole-projected raw star* additionally sums original-head predictions for reversions introduced independently into *B*, then takes half the forward sum minus half the reverse sum; (iii) *component/whole projection* first antisymmetrizes each single-substitution prediction in both endpoint sums, retaining the published hidden biases, then applies the same whole-variant projection; (iv) *JanusDDG + EvoPhi-DDG* uses construction (iii) with the MSAesm_ddG hidden biases set to zero. Each sum has a fixed starting background. Whole-variant projection is applied to the MSA-derived score before its combination with JanusDDG. Rows (ii)–(iii) isolate single-substitution projection, and rows (iii)–(iv) isolate hidden-bias removal (Methods, Sections 3.4 and 3.3). Bold marks the best displayed value per dataset and metric, including ties, and each method attaining a best value.

| Fixed predictor | Pearson | MAE | RMSE | Reversal MAE | ROC-AUC |
|---|---|---|---|---|---|
| *S669* |  |  |  |  |  |
| **JanusDDG + original MSAesm_ddG** | **0.59** | 0.92 | 1.34 | 0.10 | **0.83** |
| **JanusDDG + antisymmetric MSAesm_ddG** | **0.59** | **0.91** | **1.33** | **0.00** | **0.83** |
| **JanusDDG + EvoPhi-DDG** | **0.59** | **0.91** | **1.33** | **0.00** | **0.83** |
| *S461* |  |  |  |  |  |
| **JanusDDG + original MSAesm_ddG** | **0.75** | **0.65** | **0.87** | 0.10 | **0.93** |
| **JanusDDG + antisymmetric MSAesm_ddG** | **0.75** | **0.65** | 0.88 | **0.00** | 0.92 |
| **JanusDDG + EvoPhi-DDG** | **0.75** | **0.65** | 0.88 | **0.00** | 0.92 |
| *S96* |  |  |  |  |  |
| **JanusDDG + original MSAesm_ddG** | **0.55** | **1.45** | 2.06 | 0.12 | **0.76** |
| **JanusDDG + antisymmetric MSAesm_ddG** | **0.55** | **1.45** | **2.04** | **0.00** | 0.75 |
| **JanusDDG + EvoPhi-DDG** | **0.55** | **1.45** | **2.04** | **0.00** | 0.75 |
| *PTmul-NR* |  |  |  |  |  |
| **Janus + directional WT star** | **0.66** | **1.46** | 1.96 | 0.28 | **0.86** |
| **Janus + whole-projected raw star** | **0.66** | 1.47 | 1.96 | **0.00** | 0.85 |
| **Janus + component/whole projection** | **0.66** | 1.47 | **1.95** | **0.00** | 0.85 |
| **JanusDDG + EvoPhi-DDG** | **0.66** | 1.47 | **1.95** | **0.00** | 0.85 |

**Table S7:** Same-site transitivity on S669. Mean over ten repetitions; *N* is the number of intermediate residues. RMSE is in kcal/mol. For each *N* , bold marks the highest correlation and lowest closure RMSE across models, together with the corresponding model headings.

| *N* | Predictor | Pearson | Closure MAE | Closure RMSE |
|---|---|---|---|---|
| 1 | MSAesm_ddG original | 0.96 | 0.15 | 0.26 |
| 1 | MSAesm_ddG antisymmetrized | 1 − 3.65 × 10<sup>−7</sup> | 5.89 × 10<sup>−4</sup> | 7.78 × 10<sup>−4</sup> |
| 1 | **EvoPhi-DDG** | **1.00** | **4.26 × 10<sup>−17</sup>** | **1.03 × 10<sup>−16</sup>** |
| 1 | JanusDDG | 0.98 | 0.14 | 0.21 |
| 1 | Equal-weight ensemble | 0.99 | 0.07 | 0.11 |
| 3 | MSAesm_ddG original | 0.93 | 0.23 | 0.33 |
| 3 | MSAesm_ddG antisymmetrized | 1 − 6.11 × 10<sup>−7</sup> | 7.83 × 10<sup>−4</sup> | 1.00 × 10<sup>−3</sup> |
| 3 | **EvoPhi-DDG** | **1.00** | **7.35 × 10<sup>−17</sup>** | **1.41 × 10<sup>−16</sup>** |
| 3 | JanusDDG | 0.96 | 0.18 | 0.26 |
| 3 | Equal-weight ensemble | 0.99 | 0.09 | 0.13 |
| 5 | MSAesm_ddG original | 0.90 | 0.31 | 0.43 |
| 5 | MSAesm_ddG antisymmetrized | 1 − 8.29 × 10<sup>−7</sup> | 9.28 × 10<sup>−4</sup> | 1.17 × 10<sup>−3</sup> |
| 5 | **EvoPhi-DDG** | **1.00** | **9.62 × 10<sup>−17</sup>** | **1.74 × 10<sup>−16</sup>** |
| 5 | JanusDDG | 0.95 | 0.20 | 0.30 |
| 5 | Equal-weight ensemble | 0.99 | 0.10 | 0.15 |
| 7 | MSAesm_ddG original | 0.88 | 0.38 | 0.52 |
| 7 | MSAesm_ddG antisymmetrized | 1 − 1.07 × 10<sup>−6</sup> | 1.05 × 10<sup>−3</sup> | 1.32 × 10<sup>−3</sup> |
| 7 | **EvoPhi-DDG** | **1.00** | **1.03 × 10<sup>−16</sup>** | **1.83 × 10<sup>−16</sup>** |
| 7 | JanusDDG | 0.95 | 0.23 | 0.32 |
| 7 | Equal-weight ensemble | 0.98 | 0.11 | 0.16 |
| 9 | MSAesm_ddG original | 0.84 | 0.46 | 0.61 |
| 9 | MSAesm_ddG antisymmetrized | 1 − 1.38 × 10<sup>−6</sup> | 1.19 × 10<sup>−3</sup> | 1.50 × 10<sup>−3</sup> |
| 9 | **EvoPhi-DDG** | **1.00** | **1.22 × 10<sup>−16</sup>** | **2.07 × 10<sup>−16</sup>** |
| 9 | JanusDDG | 0.94 | 0.25 | 0.35 |
| 9 | Equal-weight ensemble | 0.98 | 0.12 | 0.17 |

**Table S6:** Paired protein-bootstrap differences in Pearson. Within each dataset, bold marks the largest ensemble-minus-comparator point estimate and its associated comparator.

| Dataset | Comparator | Δ*r* | 95% interval |
|---|---|---|---|
| S669 | JanusDDG | +0.05 | [+0.02, +0.08] |
| S669 | **EvoPhi-DDG** | **+0.06** | [+0.03, +0.09] |
| S461 | JanusDDG | +0.06 | [+0.03, +0.10] |
| S461 | **EvoPhi-DDG** | **+0.09** | [+0.05, +0.11] |
| S96 | JanusDDG | +0.04 | [-0.06, +0.15] |
| S96 | **EvoPhi-DDG** | **+0.07** | [-0.07, +0.20] |
| PTmul-NR | **JanusDDG** | **+0.05** | [+4.28 × 10<sup>−3</sup>, +0.08] |
| PTmul-NR | EvoPhi-DDG | −1.54 × 10<sup>−3</sup> | [-0.05, +0.04] |

**Table S8:**
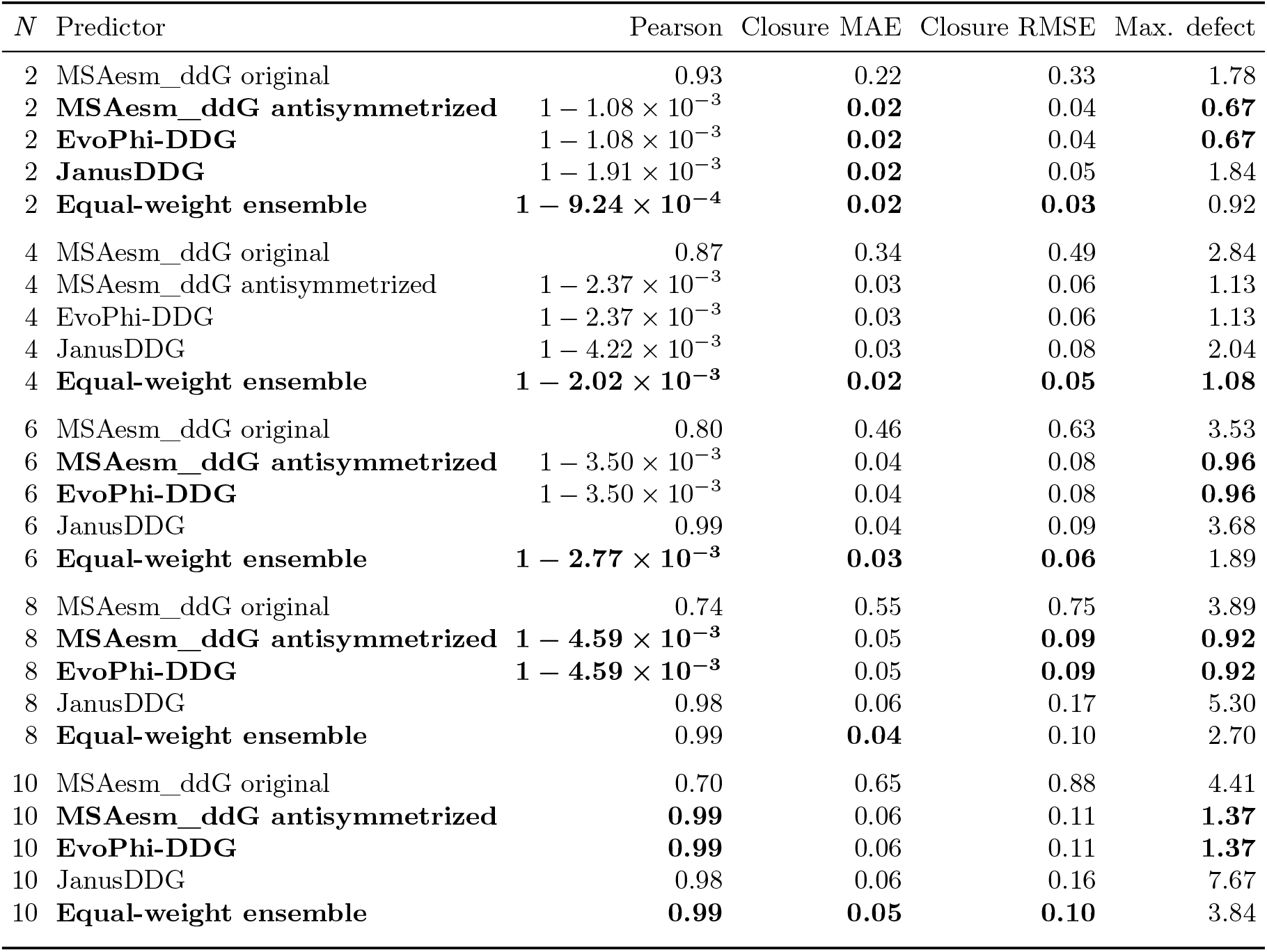
Multisite pathway consistency on S669. *N* is the number of intermediate sequence states; each path has *N* + 1 single-substitution edges. Pearson correlation and closure MAE/RMSE are means over ten repetitions of all 669 variants. Maximum absolute defect is taken over the 6,690 paths at each *N* . Errors are in kcal/mol. Each path introduces *N/*2 temporary substitutions at distinct non-target sites, applies the target substitution, and reverts temporary changes in reverse order. No experimental labels enter these metrics. Bold marks the highest correlation or lowest error for each *N* and its associated method, including ties at the displayed precision.

| *N* | Predictor | Pearson | Closure MAE | Closure RMSE | Max. defect |
|---|---|---|---|---|---|
| 2 | MSAesm_ddG original | 0.93 | 0.22 | 0.33 | 1.78 |
| 2 | **MSAesm_ddG antisymmetrized** | 1 − 1.08 × 10<sup>−3</sup> | **0.02** | 0.04 | **0.67** |
| 2 | **EvoPhi-DDG** | 1 − 1.08 × 10<sup>−3</sup> | **0.02** | 0.04 | **0.67** |
| 2 | **JanusDDG** | 1 − 1.91 × 10<sup>−3</sup> | **0.02** | 0.05 | 1.84 |
| 2 | **Equal-weight ensemble** | **1 − 9.24 × 10<sup>−4</sup>** | **0.02** | **0.03** | 0.92 |
| 4 | MSAesm_ddG original | 0.87 | 0.34 | 0.49 | 2.84 |
| 4 | MSAesm_ddG antisymmetrized | 1 − 2.37 × 10<sup>−3</sup> | 0.03 | 0.06 | 1.13 |
| 4 | EvoPhi-DDG | 1 − 2.37 × 10<sup>−3</sup> | 0.03 | 0.06 | 1.13 |
| 4 | JanusDDG | 1 − 4.22 × 10<sup>−3</sup> | 0.03 | 0.08 | 2.04 |
| 4 | **Equal-weight ensemble** | **1 − 2.02 × 10<sup>−3</sup>** | **0.02** | **0.05** | **1.08** |
| 6 | MSAesm_ddG original | 0.80 | 0.46 | 0.63 | 3.53 |
| 6 | **MSAesm_ddG antisymmetrized** | 1 − 3.50 × 10<sup>−3</sup> | 0.04 | 0.08 | **0.96** |
| 6 | **EvoPhi-DDG** | 1 − 3.50 × 10<sup>−3</sup> | 0.04 | 0.08 | **0.96** |
| 6 | JanusDDG | 0.99 | 0.04 | 0.09 | 3.68 |
| 6 | **Equal-weight ensemble** | **1 − 2.77 × 10<sup>−3</sup>** | **0.03** | **0.06** | 1.89 |
| 8 | MSAesm_ddG original | 0.74 | 0.55 | 0.75 | 3.89 |
| 8 | **MSAesm_ddG antisymmetrized** | **1 − 4.59 × 10<sup>−3</sup>** | 0.05 | **0.09** | **0.92** |
| 8 | **EvoPhi-DDG** | **1 − 4.59 × 10<sup>−3</sup>** | 0.05 | **0.09** | **0.92** |
| 8 | JanusDDG | 0.98 | 0.06 | 0.17 | 5.30 |
| 8 | **Equal-weight ensemble** | 0.99 | **0.04** | 0.10 | 2.70 |
| 10 | MSAesm_ddG original | 0.70 | 0.65 | 0.88 | 4.41 |
| 10 | **MSAesm_ddG antisymmetrized** | **0.99** | 0.06 | 0.11 | **1.37** |
| 10 | **EvoPhi-DDG** | **0.99** | 0.06 | 0.11 | **1.37** |
| 10 | JanusDDG | 0.98 | 0.06 | 0.16 | 7.67 |
| 10 | **Equal-weight ensemble** | **0.99** | **0.05** | **0.10** | 3.84 |

**Table S9:** Predictive performance across single- and multiple-substitution benchmarks. The ensemble is the fixed equal-weight mean of JanusDDG and EvoPhi-DDG. PTmul-NR uses the background-aware, whole-variant-antisymmetric EvoPhi-DDG construction. MAE and RMSE are in kcal/mol. Bold marks the highest correlation or lowest error within each benchmark and the associated methods, including ties at two decimals. Detailed comparisons are provided in Supplementary Tables S1–S4.

| Predictor | Pearson | Spearman | MAE | RMSE |
|---|---|---|---|---|
| **S669** (*n* = 669) |  |  |  |  |
| JanusDDG | 0.55 | 0.55 | 0.97 | 1.39 |
| EvoPhi-DDG | 0.53 | 0.55 | 0.99 | 1.39 |
| **Equal-weight ensemble** | **0.59** | **0.61** | **0.91** | **1.33** |
| **S461** (*n* = 461) |  |  |  |  |
| JanusDDG | 0.69 | 0.66 | 0.71 | 0.96 |
| EvoPhi-DDG | 0.66 | 0.65 | 0.73 | 0.97 |
| **Equal-weight ensemble** | **0.75** | **0.72** | **0.65** | **0.88** |
| **S96** (*n* = 96) |  |  |  |  |
| JanusDDG | 0.52 | 0.50 | 1.50 | 2.10 |
| EvoPhi-DDG | 0.48 | 0.47 | 1.53 | 2.14 |
| **Equal-weight ensemble** | **0.55** | **0.52** | **1.45** | **2.04** |
| **PTmul-NR** (*n* = 82) |  |  |  |  |
| JanusDDG | 0.61 | 0.57 | 1.57 | 2.06 |
| **EvoPhi-DDG** | **0.66** | 0.58 | **1.45** | **1.95** |
| **Equal-weight ensemble** | **0.66** | **0.61** | 1.47 | **1.95** |

**Table S10:**
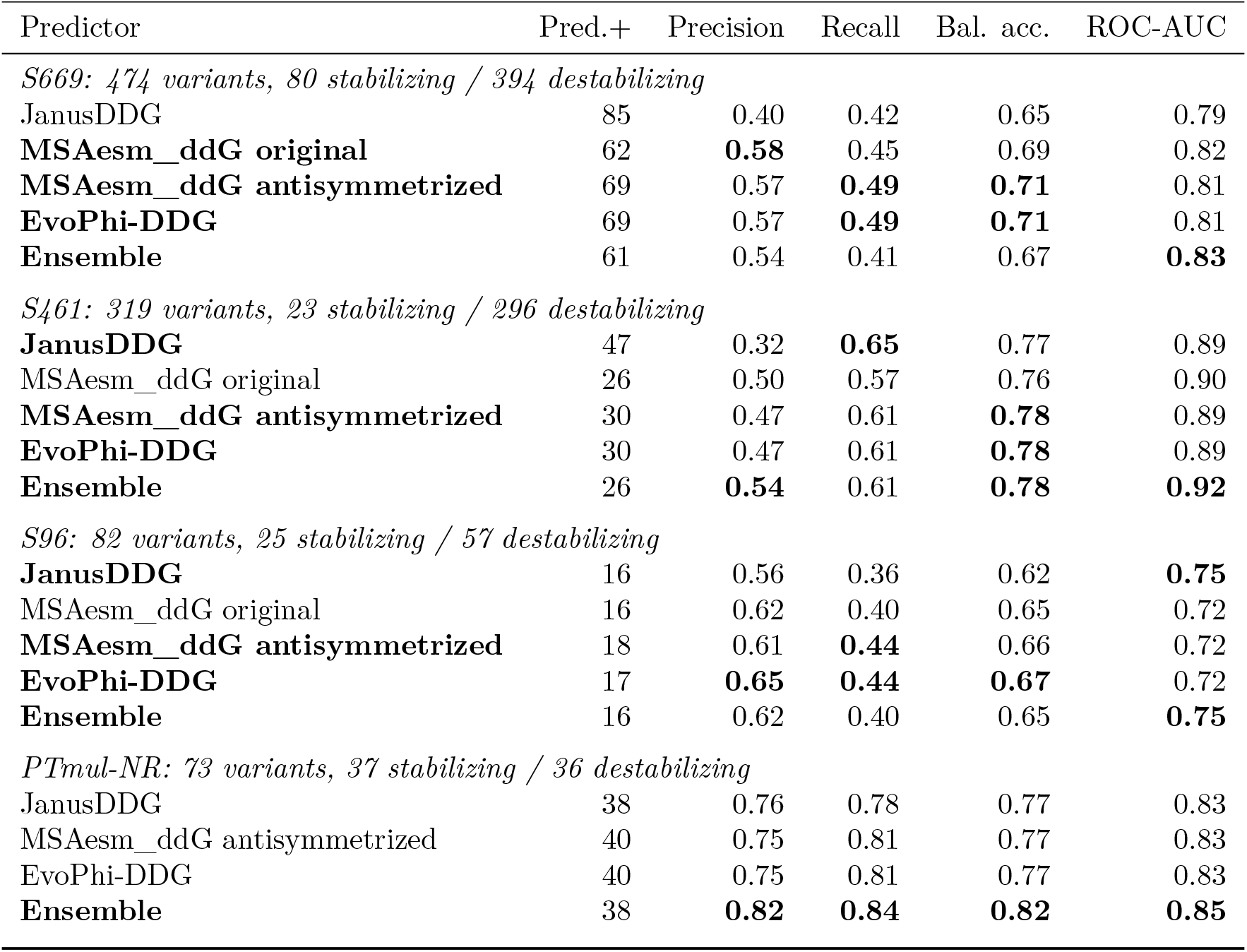
Classification after excluding experimentally near-neutral variants. Positive means stabilizing. Pred.+ is the number predicted stabilizing using the fixed zero score threshold. Precision and recall refer to the stabilizing class; balanced accuracy averages the two class recalls. ROC-AUC uses continuous scores. The same experimental mask is applied to all models within a dataset. Best metric values and associated methods are bold within each dataset, including ties at the displayed precision. Pred.+ is a descriptive count and is not ranked.

